# Extreme conservation of cnidarian stinging cell identity despite 600 million years of evolution

**DOI:** 10.64898/2026.06.16.732747

**Authors:** Sarah E Arnold, Ryan M Besemer, Koty Sharp, Loretta M Roberson, Jacob F Warner, Leslie S Babonis

**Affiliations:** Department of Ecology and Evolutionary Biology, Cornell University, Ithaca NY USA; Department of Biology and Marine Biology, UNC-Wilmington, Wilmington NC USA; Department of Biology, Marine Biology, and Environmental Science, Roger Williams University, Bristol RI USA; Marine Biological Laboratory, Bell Center, Woods Hole MA USA

## Abstract

Understanding how cells specialize is essential for reconstructing the diversification of life on earth. Cnidocytes (stinging cells) have a single origin in the stem cnidarian (∼800mya) and have since specialized into extremes in morphology and function. Using single-cell RNA sequencing and transcriptional lineage reconstruction in a coral and a sea anemone, we show that a single gene (FoxL2) controls a critical switch point in the evolution of cnidocyte diversity: the decision to be a piercing cell (nematocyte) or an ensnaring cell (spirocyte). Nematocytes are lowly conserved between species and express younger genes, reflecting the continued diversification of this cell lineage. Surprisingly, ensnaring cells are one of the most highly conserved differentiated cell types and express older genes than nematocytes. This suggests spirocytes reached an adaptive peak early and have changed little during the 600 million years since corals and sea anemones last shared a common ancestor, making them ‘living fossil’ cells.

## Introduction

Animals evolved from a unicellular ancestor; thus, the astonishing variety of animal forms and functions present on the planet is a result of extraordinary cell type diversification. Without access to the cells of our animal ancestors, efforts to understand how cell identity evolves rely largely on comparisons of homologous cells from distantly related taxa^1–5^. The advent of single-cell RNA sequencing (scRNA-seq) has revolutionized our ability to compare cell types across species to understand cell type diversification^6^. These comparisons have proven difficult, however, as animal cell types have diversified so greatly over the past 850 million years of evolution from our last common ancestor^7^ and it is not always obvious which cell types are homologous when comparing two animal taxa. Additionally, models of how much the gene expression in homologous cells should vary over different evolutionary distances are lacking^6^.

Cnidocytes, the infamous stinging cells found only in cnidarians (jellyfish, hydroids, sea anemones, and corals), provide an unparalleled opportunity to compare homologous cell types across deep evolutionary timescales. These novel cells arose once in the last common ancestor of cnidarians^8^ and have since diversified into a clade of cells with extraordinary variation in form and function^9,10^. All cnidocytes contain a cnidocyst, the organelle responsible for the sting, which is composed of a pressurized capsule containing an eversible tubule^11^. Upon triggering, the tubule is discharged nearly instantaneously in one of the fastest naturally occurring cellular processes^12^. In nematocytes (piercing cells) the tubule is typically adorned with spines and loaded with venom, used to penetrate and immobilize prey^10^. Nematocytes are found in both anthozoans (sea anemones, hard corals, soft corals, and tube anemones) and medusozoans (true jellyfish, box jellies, stalked jellies, and hydroids), and these cells exhibit considerable diversity in the size, shape, and ornamentation of their tubule as well as in the type of venom they contain. As many as eight morphologically distinct nematocytes have been described from a single organism^13^ and different morphological types are thought to play unique roles in prey capture^10,14,15^. Spirocytes are a non-nematocyte cnidocyte found only in hexacorals (sea anemones, hard corals, and tube anemones), making this an important clade for characterizing the mechanisms driving cnidocyte evolution^16^. They do not contain venom or pierce prey; when triggered, they release tubules adorned with sticky microfibrils to ensnare prey^9^. Despite 700 million years of evolution resulting in over 4000 extant species of hexacorals^17^, spirocytes have diversified into only two morphological types: robust and gracile^18^. This conservation is a stark contrast with the tremendous variation exhibited by nematocytes, despite both cell types evolving for over half a billion years. Here, we investigate how homologous cell types diversify over large evolutionary distances by comparing cnidocytes from the model sea anemone, *Nematostella vectensis*, with cnidocytes from an emerging model coral, *Astrangia poculata.* We present the first comprehensive single-cell RNA-seq atlas for early coral development and use this resource to examine the age and conservation of genes in homologous cell types from the two taxa. With this analysis, we demonstrate that nematocytes exhibit low conservation and appear to be evolving under diversifying selection whereas spirocytes are indistinguishable in these two taxa, suggesting these cells are evolving under strong purifying selection. These data provide an important framework for linking the evolutionary forces that drive cell diversity with the gene expression changes underlying functional diversification.

## Results

### Identifying the embryonic cell types present in the last common ancestor of sea anemones and corals

Comparative analysis of embryogenesis in *Astrangia poculata* and *Nematostella vectensis* provides valuable insight into the evolution of cell type differentiation over a large evolutionary distance. Like *N. vectensis*, *A. poculata* embryos are asynchronous in early development, such that at 12 hours post fertilization (hpf) and 24hpf we observed embryos with varying morphologies (**Fig 1A**). Gastrulation begins at 24hpf via invagination as actin-rich pseudopodia extend into the blastocoel from the presumptive endodermal cells and attach to the basal side of the presumptive ectodermal cells, similar to what has been described previously for *N. vectensis*^19,20^. These presumptive endodermal cells are ciliated in *A. poculata*, with the cilia projecting into the blastocoel (**Fig. 1A** – late blastula inset). By the early planula stage (36 and 48hpf), gastrulation is complete, and the pharynx has begun to invaginate. This stage is also typified by the presence of an apical tuft, previously described in *A. poculata* but uncommon among corals^21,22^. Despite similarities in early embryogenesis, later stages of development show striking differences between *A. poculata* (**Fig. 1A**) and *N. vectensis* (**Fig. 1B**). *N. vectensis* has a short, non-feeding larval stage during which they make their mesenteries (multifunctional digestive/reproductive tissues) by secondary invagination of the oral ectoderm^23^. The larval stage is followed by metamorphosis into a polyp, which occurs without a cue within a week after fertilization in *N. vectensis*^24^. By contrast, *A. poculata* does not make mesenteries as larvae; development pauses after invagination of the pharynx at the late planula stage despite feeding (**Fig. 1A, B**) and metamorphosis into a polyp is only initiated after a suitable settlement cue has been received^22^.

**Figure 1:**
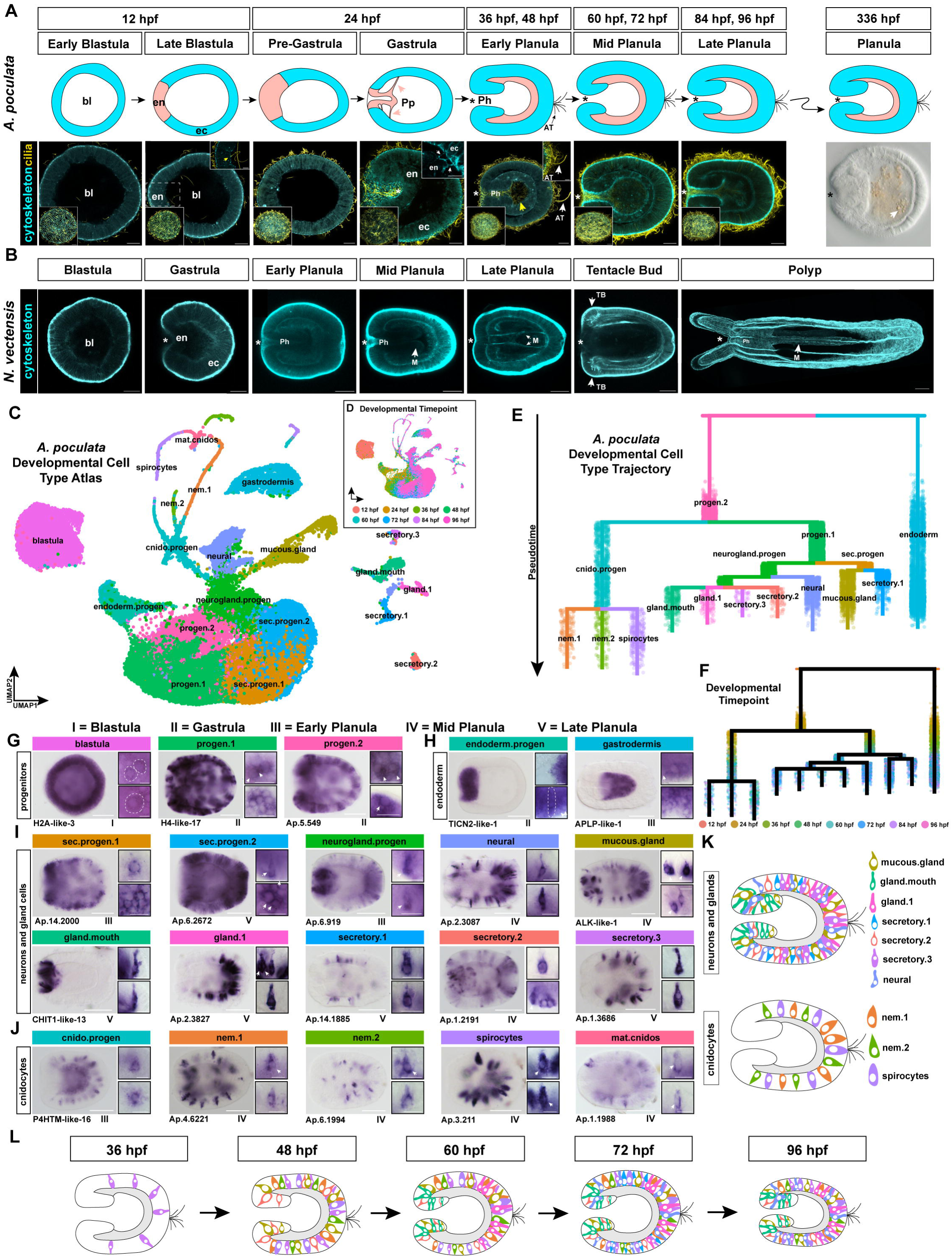
Analysis of cell differentiation during early development in *Astrangia poculata* reveals 20 cell populations with distinct gene expression patterns. **A)** Overview of morphological changes during early development. F-actin (cytoskeleton) labeled in blue and α-acetylated tubulin (cilia) labeled in yellow. Main images are max projections of a few slices at the midpoint of the embryo and lower insets are max projections showing a surface view. Yellow arrows indicate ciliated endoderm. Late blastula upper inset shows ciliated endoderm progenitors. Gastrula upper inset shows invaginating endodermal cells with pseudopodia. Early planula upper inset shows apical tuft. DIC image of live embryo at 336hpf; white arrow indicates food in the planula gut. Scale bars are 20μm for whole embryo images and 10μm for insets. en = endoderm (pink), ec = ectoderm (blue), AT = apical tuft, Ph = pharynx, Pp = pseudopodia, * = mouth. **B)** Overview of morphological changes during the first week of development in *Nematostella vectensis,* as previously described^19,20^. F-actin (cytoskeleton) labeled in blue. Scale bars are 50μm. TB = tentacle bud, M = mesenteries. **C)** Merged *A. poculata* developmental UMAP with color indicating cell type. **D)** The same UMAP with color indicating developmental timepoint. **E)** *A. poculata* developmental cell type trajectory tree generated with URD. Colors match the corresponding cell populations in C. **F)** The same URD tree with color indicating developmental timepoint. **G-J)** In situ hybridization of cell type marker expression in whole embryos; side panels show cell morphologies in higher magnification; roman numerals indicate developmental stage of image. Colors as indicated in C. Scale bars are 50μm for whole embryo images and 10μm for side panels. Representative expression patterns are shown for (G) three populations of early progenitor cells; (H) two endodermal cell populations (differentiating progenitors and gastrodermis); (I) neuron + gland cell populations including: two populations of secretory progenitor cells, a population of neuroglandular progenitors, one neural cell type, and six secretory cell populations; and (J) five cnidocyte populations. **K)** Schematics summarizing the distributions of neuroglandular cells and cnidocytes throughout the embryonic ectoderm (endoderm in grey). **L)** Schematic summarizing the onset of neuroglandular cell differentiation. Cell colors as indicated in K.

The development of single-cell sequencing technologies has allowed for the generation of numerous cell atlases from across cnidarians^2,25–31^. These resources enable unparalleled study of differentiated cell types, but many of these atlases are restricted to adult tissues or single developmental timepoints and cannot be used to elucidate the early decisions underlying cell type specification. Atlases capturing stem/progenitor cell differentiation have been generated during embryonic development in *Nematostella vectensis*^2,25,26^ and during adult tissue homeostasis in the medusozoan *Hydra vulgaris*^27^, revealing a common origin of the major ectodermal cell types: cnidocytes, neurons, and gland cells. To investigate how gene expression drives cell specification and how conserved cell differentiation trajectories vary during embryogenesis, we dissociated *A. poculata* embryos every 12 hours up to 96hpf and performed scRNA-seq (**Fig. 1C**). We found 20 cell populations with distinct gene expression patterns and identified major larval cell types previously described for *N. vectensis* (neurons, gland cells, cnidocytes, and gastrodermis; **Fig. 1C, D**). To reconstruct the trajectory of cell type differentiation, we generated a developmental lineage tree using URD^32^ (**Fig. 1E, F**). We found that cnidocytes, neurons, and gland cells originate from a common pool of progenitors, recapitulating the close developmental relationship between neurons and gland cells as has been previously shown^25,27^. We also observed that some secretory cell populations form a clade with neurons whereas others differentiate from an independent, secretory progenitor. This suggests there may be greater diversity in the evolutionary origin of cnidarian secretory cells than previously appreciated. Genes expressed highly (>2logFC) and exclusively in each cluster were chosen as markers for each cell population and were verified by in situ hybridization (**Fig. 1G-J**), showing the broad distribution of early progenitors and scattered distribution of differentiated cell types (**Fig. 1K)**. By projecting progenitor marker genes onto the URD tree, we identified the progenitor populations that lead to specific differentiated cell populations (**Fig. S1**). Previous studies of *N. vectensis* have shown that cell differentiation occurs early and continuously during embryogenesis, with genes expressed in terminally differentiating cell types (e.g., neurons and cnidocytes) first appearing in the presumptive ectoderm in a salt and pepper pattern at gastrulation^33,34^. Differentiated cell populations do not appear until the early planula stage, later than is observed in *N. vectensis* (**Fig. 1D, F, L**).

### Cnidocytes and secretory cells exhibit extreme conservation, neurons do not

To understand how homologous cell identities have diverged over the 600 million years since *A. poculata* and *N. vectensis* last shared a common ancestor (**Fig. 2A**), we compared cells in the neuroglandular lineage (neurons, gland cells, and cnidocytes) using SAMap^3^. We saw expected alignment between broad homologous cell categories but observed inconsistency in the strength of alignment of different cell categories, varying from very high (mature cnidocytes; alignment score 0.85) to low (neuron and gland cell progenitors; alignment score 0.25). (**Fig. 2B**). We next investigated if subpopulations of these ectodermal cells (e.g. broad category: secretory cells, subpopulation: mouth gland cells) are conserved. We extracted cnidocytes, neurons, and gland cells (**Fig. 2C**, **Fig. S2**, **Fig. 3A**) from the full *A. poculata* scRNA-seq dataset, re-clustered them to generate cell type specific subsets, and compared these subsets with *N. vectensis*^25,26^. We saw strikingly high alignment (clear 1:1 alignment with scores > 0.5) in some populations of cnidocytes and secretory cells, indicating clear homology and strong conservation of these cell types. In contrast, neural subpopulations exhibited low alignment (few 1:1 alignments, majority < 0.1) between species, recapitulating what has been shown in teleosts and broadly across bilaterians^4,5^. GLWamide+ and RFamide+ neurons are two of the most well studied neural subpopulations in cnidarians^35^, so we investigated their conservation between *A. poculata* and *N. vectensis* larvae. Despite shared neuropeptide expression, these neural subpopulations do not align as highly as secretory cells and cnidocytes between species (GLWamide+ neuron alignment scores: 0.22, 0.03, 0.04, 0.02; RFamide+ neuron alignment scores: 0.26, 0.09; **Fig. S2B-D**). Consistent with this, comparison of progenitor cells in *A. poculata* and *N. vectensis* revealed surprisingly weak alignment for shared neuron and gland cell progenitors. We hypothesize this may be attributed to the heterogeneous nature of these progenitors as they give rise to numerous diverse neural and gland subpopulations in cnidarians^27,36^. Cell types expressing a greater number of genes have more potential for changes in gene expression and therefore might exhibit lower alignment between species, but we found no relationship between number of genes expressed per cluster and alignment score (**Fig. S3A, B**). To determine if expression of genes associated with particular cellular functions drives alignment between cell types, we conducted functional enrichment analysis using SAMap. We found that many cnidocyte and secretory clusters were enriched for genes involved in translation and ribosome biogenesis as these cells make large quantities of the proteins they secrete; however, there are many secretory and cnidocyte cell clusters enriched for this category that do not have high alignment between species, so it is unlikely that this functional enrichment is driving alignment (**Fig S3C**).

**Figure 2:**
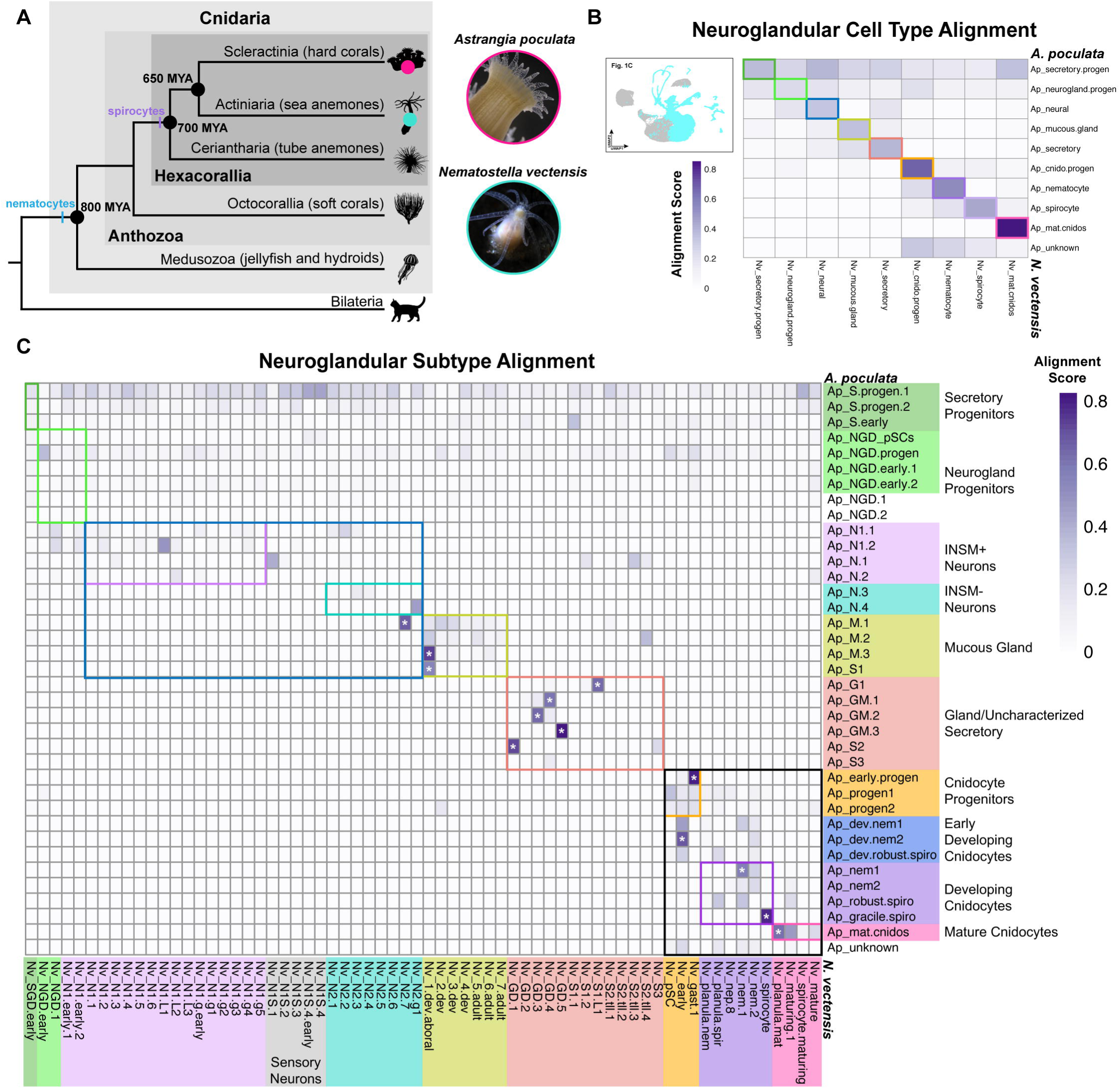
Comparison of ectodermal cell types in *A. poculata* and *N. vectensis* reveals surprisingly variable degrees of transcriptomic conservation. **A)** Tree showing placement of *A. poculata* and *N. vectensis*. Nematocytes evolved in the last common ancestor of all cnidarians, while spirocytes evolved in the last common ancestor of hexacorals. Node ages as reported in McFadden et al., 2022^17^. **B)** Alignment of neuroglandular cell categories from *A. poculata* (**Fig 1C**) and *N. vectensis*^25,26^ represented as a heat map. Alignment scores generated in SAMap. **C)** Heat map of neuroglandular subpopulation alignment scores. Colored boxes correspond to cell categories in C. NGD = neuroglandular, N = neural, M = mucous, S = secretory, and GM = gland mouth. * = alignment scores greater than 0.5.

**Figure 3:**
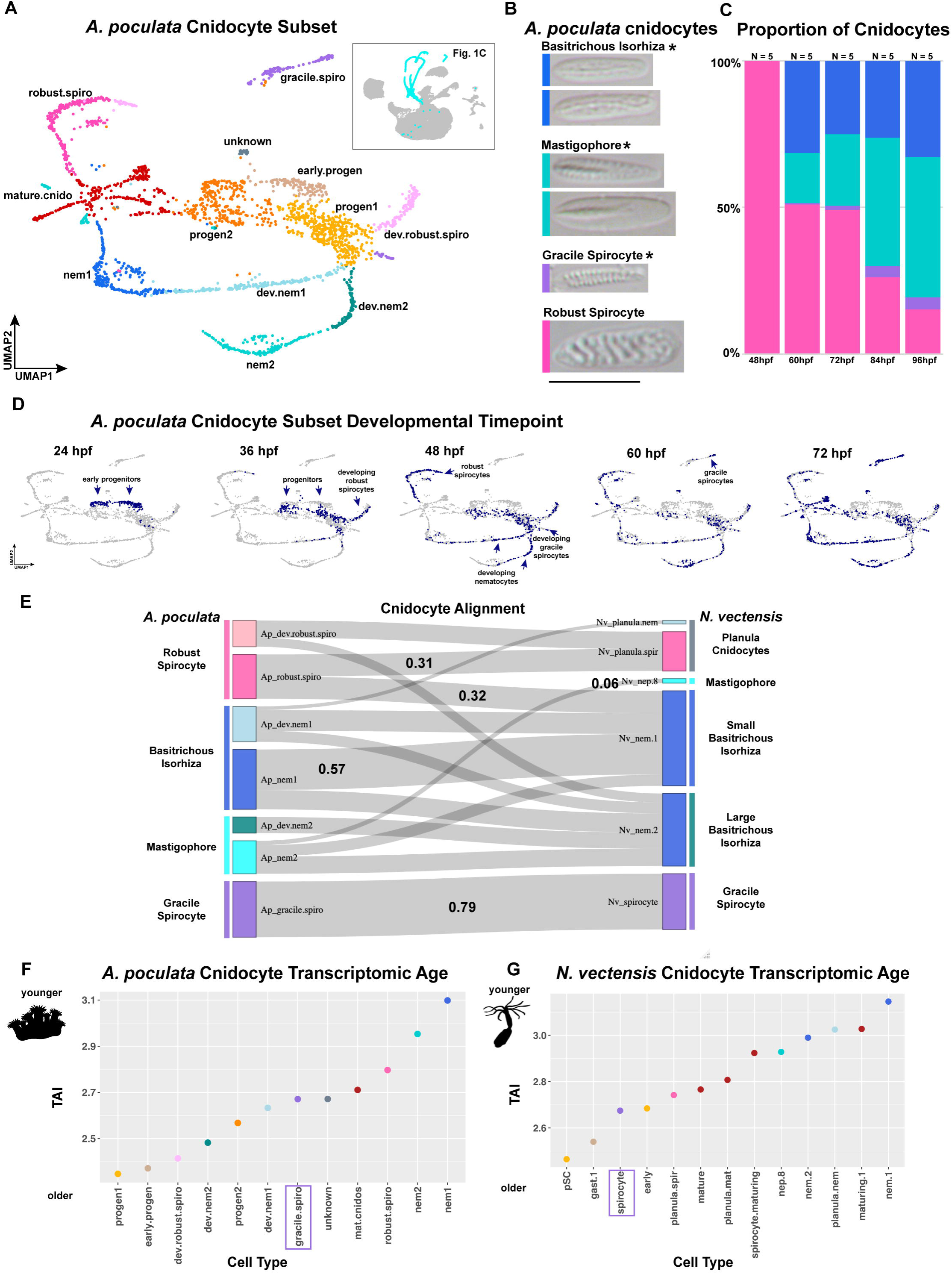
Characterization of cnidocyte development in *Astrangia poculata*. **A)** UMAP of re-clustered cnidocytes from Figure 1C annotated using known cnidocyte markers from *N. vectensis*. **B)** Four types of cnidocyte arise during the first 96 hours of embryogenesis, shown with DIC microscopy. Scale bar is 10μm. * indicate cnidocyte types that are also present in *N. vectensis*. **C)** The proportion of each cnidocyte type present varies with developmental stage. No mature cnidocytes were detected before 48hpf. N = number of embryos analyzed per timepoint. **D)** Cnidocyte subset UMAPs with color (navy blue) indicating developmental timepoint. **E)** Sankey plot of different cnidocyte types in *A. poculata* and *N. vectensis* with flows between cell types representing alignment scores. Colors of *A. poculata* cnidocyte clusters correspond to the UMAP in A. Colors of *N. vectensis* cnidocyte clusters correspond to the *A. poculata* cnidocyte cluster they most highly align with. Thicker flows between cells represent higher alignment. Some *A. poculata* cnidocytes align with multiple *N. vectensis* cnidocyte types, but gracile spirocytes are the only cells that exclusively align with one another between species. Numbers on flows indicate alignment scores discussed in text. **F)** Analysis of gene age in each cnidocyte cluster in *A. poculata*. Y-axis shows transcription age index (TAI); larger TAI values correspond to younger gene ages. Dot color corresponds to cell type color in **A**. **G)** Analysis of gene age in each cnidocyte cluster in *N. vectensis*. Dot colors for *N. vectensis* correspond to the *A. poculata* cnidocyte cluster with the highest alignment.

We hypothesized that homologous cell types with low alignment between species may express a greater number of “young” or lineage-restricted genes as this could account for the high transcriptomic variation. To test this, we performed a phylostratigraphy analysis following the methods of Piovani et al. (2023)^37^ using the programs GenEra^38^ and myTAI^39^ to estimate the transcriptome age index (TAI) for each cell cluster in *A. poculata* embryos (**Fig. S3D**). Similar to previous work, we found that generally, progenitor cells have older transcriptomic age than differentiated cells, likely reflecting the conservation of genes that control the maintenance of pluripotency and proliferation^37^. We also found that cnidocyte clusters have the youngest transcriptomic age (highest TAI), reflecting the role of taxonomically restricted genes such as minicollagens (cnidocyte structural components)^40^ in driving the evolution of this cnidarian novelty. We also found that neurons are transcriptomically younger than secretory cells in *A. poculata*. If neurons are expressing and incorporating novel genes into their gene regulatory networks at greater levels than secretory cells, this may drive the observed low alignment of neural populations and high alignment of secretory cell populations between species.

### Spirocyte transcriptional identity is highly conserved

Overall, cells in the cnidocyte lineage exhibited the greatest variation in alignment across species, suggesting distinct evolutionary processes may be driving the diversification of these cell types. To further elucidate the mechanisms underlying the observed variation in homologous cell type evolution, we investigated the development of cnidocytes in *A. poculata.* Studies of embryonic cnidocyte development have previously been limited to *N. vectensis,* but with access to the embryonic cell atlas from *A. poculata*, we can explicitly test conservation in cnidocyte development across species.

To generate a cnidocyte subset scRNA-seq dataset, we extracted and re-clustered cnidocytes from the full *A. poculata* scRNA-seq dataset and annotated the resulting clusters based on expression of known cnidocyte subpopulation genes in *N. vectensis* (**Fig. 3A**). We then characterized the types of cnidocytes observed and their frequency over development using morphological analysis. We found four types of cnidocytes arise during embryogenesis in *A. poculata*: two types of basitrichous isorhizas (nematocytes), two types of mastigophores (nematocytes), and both types of spirocytes (robust and gracile) (**Fig. 3B**). The morphology and distribution of three of these, basitrichous isorhizas, mastigophores, and gracile spirocytes, were described previously from adult tissues^41^. Robust spirocytes are the first cnidocyte to develop in embryos and comprise 100% of the mature cnidocytes at 48hpf (**Fig. 3C**). By 72hpf all three other types are present but in variable proportions that change over developmental time. To connect these morphological data with our scRNA-seq data, we plotted key developmental timepoints on the cnidocyte subset UMAP and show that progenitors are present at 24hpf and robust spirocytes are developing by 36hpf (**Fig. 3D**). This indicates that cnidocytes take between 12 and 24 hours to mature in *A. poculata*.

Three of the four cnidocyte types found in *A. poculata* embryos are also found in *N. vectensis* embryos (basitrichous isorhizas, mastigophores, and gracile spirocytes)^42^ (**Fig 3B, 4A**). Comparing the homologous cnidocyte types across species using SAMap revealed considerable variation in alignment scores (**Fig. 3E**). Most notably, the gracile spirocytes of both species show the highest alignment of all cnidocyte types (0.79) and are the only cells that do not share alignment with any other cnidocyte types (**Fig. 3E**). Nematocytes were highly variable: while small basitrichous isorhizas from *N. vectensis* have high alignment (0.57) with the basitrichous isorhizas from *A. poculata,* mastigophores from the two species have very low alignment (0.06). Robust spirocytes are not typically found in *N. vectensis;* however, it was recently shown that knockout of one gene (*NvSox2*) in *N. vectensis* resulted in transformation of the small basitrichous isorhizas into robust spirocytes, revealing a close (and unexpected) relationship between these two cell types^16^. In support of this result, we find the robust spirocytes from *A. poculata* to have equally high alignment (0.32) with small basitrichous isorhizas and a previously uncharacterized population of larval (planula) spirocytes (0.31).

We hypothesized that nematocytes may express more novel genes than spirocytes reflecting their continued diversification throughout anthozoans. To test this, we performed phylostratigraphy analysis ^38,39^ on the cnidocyte clusters of both species and found that nematocytes have a higher TAI (younger transcriptomic age) than the gracile spirocytes in both species (**Fig. 3F, G**). This supports previous morphological observations that gracile spirocytes have changed very little since their origin in the ancestor of hexacorals and explains the high alignment that we see between *A. poculata* and *N. vectensis* spirocytes. Interestingly, we found that small basitrichous isorhizas have the youngest transcriptomic age of all cnidocytes. This supports previous work demonstrating that this type of nematocyte evolved from a spirocyte in the lineage leading to *N. vectensis* ^16^ and suggests the basitrichous isorhiza morphology may have evolved multiple times from different suites of genes. We also found that mature cnidocytes have an older transcriptomic age than developing cnidocytes, suggesting that the physiology of the cell housing the cnidocyst may not be changing very much over time while the genes specifying the size and shape of the cnidocyst continue to evolve.

Given the strong alignment of some cnidocyte types, we sought to investigate whether genes previously identified in the cnidocyte gene regulatory network in *N. vectensis* are conserved in *A. poculata*. These include the transcription factors *NvSoxB2*^34^, *NvCnZnf1*^43^, *NvPaxA*^44^, and *NvSox2*^16^ and several cnidocyte-specific structural molecules called minicollagens^42^ (**Fig. 4B**). Both the upstream transcription factors and the downstream effectors (minicollagens) are expressed in developing cnidocytes in *A. poculata* at embryonic stages consistent with their sequential activation during cnidocyte differentiation (**Fig. 4B-G**). This confirms conservation of the core cnidocyte GRN in the ancestor of corals and sea anemones.

**Figure 4:**
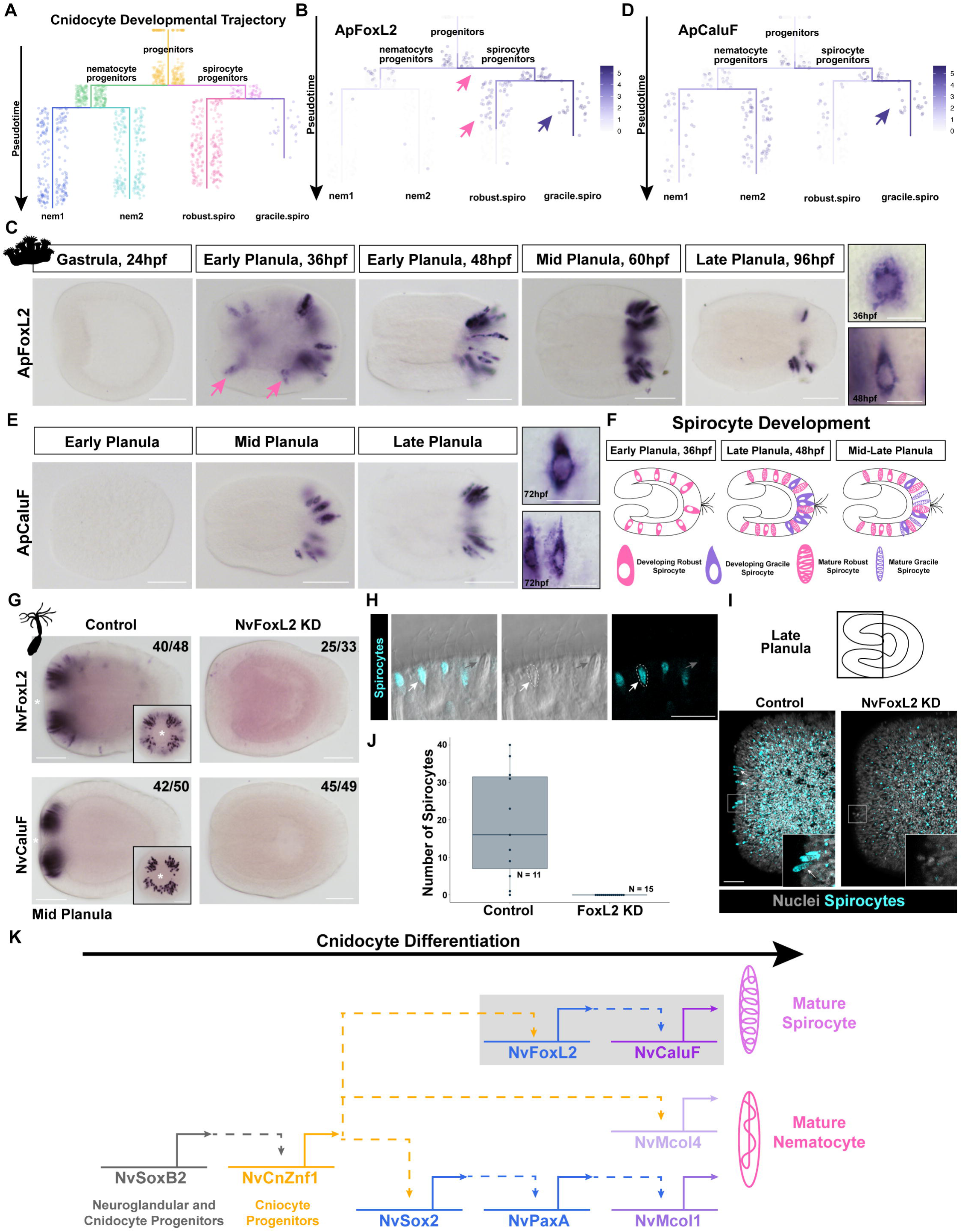
Cnidocyte gene regulatory network components are expressed in *A. poculata.* **A)** *N. vectensis* has three types of cnidocytes. Scale bar is 10μm. **B)** A gene regulatory network of known cnidocyte developmental genes in *N. vectensis* (see:^16,34,43,44^). **C)** Dot plot showing expression of known cnidocyte development genes in the scRNAseq cnidocyte subset data. Size of circle indicates the percentage of cells in each cluster expressing each gene. Color of circle indicates the normalized average expression of each gene within each cluster. **D)** In situ hybridization of known cnidocyte developmental regulators in *A. poculata*. **E)** Maximum likelihood tree of minicollagen genes from *N. vectensis* and *A. poculata.* Node labels are bootstraps. **F)** In situ hybridization of minicollagens at the mid planula stage (60hpf). **G)** Expression of each minicollagen projected onto the cnidocyte subset UMAP. **H)** Summary of the specific repertoires of minicollagens expressed in different cnidocyte types. All scale bars are 50μm.

To put this analysis of cnidocyte development in a broader phylogenetic context, we compared the cnidocyte clusters of *A. poculata* and *N. vectensis* to those of the octocoral *Xenia sp.* and the medusozoan hydroid *Hydra vulgaris,* using previously generated scRNA-seq data^27,29^ (**Fig. S4**). We observed that progenitor cells and mature cnidocytes aligned across species, suggesting conserved gene expression in these populations despite large evolutionary divergence. Surprisingly, we saw high alignment of *A. poculata* basitrichous isorhizas (nem1) and mastigophores (nem2) with *H. vulgaris* stenoteles (nb5; alignment scores of 0.66 and 0.63, respectively) despite stenoteles being Hydrozoan specific cnidocytes^45^. This suggests similar genes have been used in these distinct cell types to make taxon-specific morphologies. Future studies knocking down the orthologous genes in each species would be useful for assessing how distant taxa use the same genes to make distinct cnidocyte morphologies. We also saw high alignment of *N. vectensis* and *A. poculata* small basitrichous isorhizas with *Xenia* atrichous isorhizas (alignment scores of 0.58 and 0.64, respectively). Atrichous isorhizas are hypothesized to have evolved in the common ancestor of all cnidarians and most closely represent the ancestral cnidocyte from which all other types diversified^8^. We suggest the high alignment of atrichous isorhizas from octocorals and basitrichous isorhizas from hexacorals reflects the shared genes necessary for patterning the ground state of the isorhiza cnidocyte and we expect the genes expressed uniquely in basitrichous isorhizas are likely to encode the spines, which are lacking in atrichous isorhizas. Future studies could test this hypothesis by manipulating basitrichous isorhiza specific genes in *A. poculata* or *N. vectensis* in attempt to restore the atrichous isorhiza phenotype.

### FoxL2 is a novel regulator of cnidogenesis

Cnidocytes develop continuously and asynchronously throughout embryogenesis, enabling us to capture various stages of cnidocyte development in our cell atlas. We next wanted to reconstruct cnidocyte developmental trajectories to predict intermediate stages that cannot be visualized morphologically. We used the lineage inference trajectory program URD^32^ to construct a cnidocyte developmental trajectory from our cnidocyte scRNA-seq dataset (**Fig. 5A**) and found that nematocyte and spirocyte fates split early during differentiation. This confirms that all four cnidocyte types differentiate from a common progenitor and then progress through nematocyte and spirocyte specific progenitor states before terminally differentiating into distinct cnidocyte types (**Fig. 5A**).

**Figure 5:** FoxL2 drives spirocyte development in *A. poculata* and *N. vectensis*. **A)** A cnidocyte developmental trajectory tree generated by URD. Early progenitor cells (tan, Fig. 3A) were chosen as the root and each cnidocyte type cluster was chosen as the tips of branches. **B)** *ApFoxL2* expression mapped onto the cnidocyte URD tree shown in Fig 5A. Pink arrows indicate *ApFoxL2* expressed in shared robust and gracile spirocyte progenitors as well as early in robust spirocyte differentiation. Purple arrow indicates *ApFoxL2* expression in differentiating gracile spirocytes. **C)** In situ hybridization of *ApFoxL2*. Pink arrows indicate putative developing robust spirocytes. Side panels show cell morphologies at higher magnification; staining is excluded from the nucleus which we see here as a large empty space in the cell. **D)** *ApCaluF* expression mapped onto the cnidocyte URD tree shown in Fig 5A. **E)** In situ hybridization of *ApCaluF*; side panels show cell morphologies at higher magnification. Scale bars are 50μm for whole embryo images and 10μm for side panels. **F)** A summary of spirocyte development in *A. poculata*. **G)** In situ hybridization of *NvFoxL2* and *NvCaluF* in control and *NvFoxL2* knockdown mid planula embryos. Insets show oral view of expression in controls. Star indicates the mouth. Scale bars are 50μm. **H)** Staining with FITC tyramide labels mature spirocytes, but not nematocytes. Scale bar is 10μm. White arrows indicate spirocytes; grey arrows indicate nematocytes. **I)** Spirocytes (blue) and nuclei (grey) are shown in the tentacle bud region of late planula controls and *FoxL2* knockdown embryos. Scale bar is 20μm. **J)** Spirocytes are lost in *NvFoxL2* knockdown embryos. **K)** Gene regulatory network of cnidocyte development in *N. vectensis*. Grey box indicates data from this paper. Other data from:^16,34,43,44^.

Spirocyte development has not been characterized previously in any cnidarian. We found that the transcription factor *FoxL2* was the top expressed transcription factor in the spirocyte progenitor in *A. poculata*, and the expression of this gene is maintained during differentiation of gracile spirocytes and downregulated during late differentiation of robust spirocytes (**Fig. 5B**). Visualization of *FoxL2* using in situ hybridization shows expression in developing cnidocytes throughout the embryo at 36hpf. When robust spirocytes are maturing at 48hpf, expression of *FoxL2* in cnidocytes is restricted to the aboral region (**Fig. 5C**), consistent with the distribution of developing gracile spirocytes at this time. Rare cells expressing *FoxL2* that do not appear to be cnidocytes were also seen in the aboral region of the embryo at 48hpf (**Fig. S5A**). Based on previous reports of *FoxL2* expression in neurons in *N. vectensis*^2,46^, we refer to these cells as putative neurons; confirmation of their identity will require future investigation. To further characterize the gene regulatory network of spirocyte differentiation in *A. poculata,* we investigated the expression of *CaluF*, a gene previously described to be expressed only in the gracile spirocytes in *N. vectensis*^16^. We found that *CaluF* is expressed in gracile spirocytes but not robust spirocytes in *A. poculata* (**Fig. 5D**) and its expression is restricted to developing cnidocytes in the aboral region of the embryo (**Fig. 5E**). Therefore, the developing *FoxL2+* cnidocytes in the oral and middle regions of the embryo at 36hpf are robust spirocytes, while those in the aboral region include both robust and gracile spirocytes (**Fig. 5F**). These results reflect the morphological analyses (**Fig. 3C**) showing an increasing proportion of gracile spirocytes throughout development and a decreasing proportion of robust spirocytes. Collectively, this suggests robust spirocytes are specified in a short pulse during early development while gracile spirocytes are specified continuously.

*FoxL2* has previously been shown to be expressed GLWamide+ neurons and spirocytes in *N. vectensis*^2^ (**Fig. S5B**), but its role in their development has not been determined. To test if *FoxL2* is required for gracile spirocyte development, we knocked it down in *N. vectensis* using short hairpin RNAs^47^. We show complete loss of mature gracile spirocytes and loss of expression of the gracile spirocyte marker, *CaluF* (**Fig. 5G-J, Fig. S5C-D**). We also observed a small but significant decrease in the number of GLWamide+ neurons in mid planula embryos (p = 0.01; **Fig. S5E-F**). These data suggest FoxL2 plays a crucial, conserved function in the specification and development of spirocytes during hexacoral embryogenesis (**Fig 5K**). The role of *FoxL2* in the specification of GLWamide+ neurons appears to be less consistent and warrants future study.

## Discussion

Understanding how cell identity changes over time to facilitate organismal evolution is an ongoing challenge in evolutionary biology. The development of tools for generating and analyzing scRNA-seq data have now made it possible to compare cell identity across highly divergent taxa^48,49^. These tools identify homologous cell types based on gene expression, but how this relates to developmental homology requires fine sampling of early developmental stages in closely related taxa. In this study, we compared homologous cells (cnidocytes) in two closely related animals separated by 600 million years of evolution (corals and sea anemones) and find evidence of both extreme conservation (spirocytes) and extreme divergence (nematocytes). Cnidocytes are, therefore, an ideal model for reconstructing the diverse evolutionary forces that influence how cell identity evolves.

Previous studies of bilaterian taxa have shown that broad cell type comparisons (e.g., neurons) exhibit high alignment across taxa whereas comparisons of subtypes (e.g., GLWamide+ neurons) show lower alignment. This pattern is hypothesized to reflect the preservation of general shared cell type modules that confer functions required for cell type identity (e.g., a general neural module) and divergence in the modules that confer identity at the level of subtype^4,5^. In cnidarians, broad comparisons of neurons, secretory cells, and cnidocytes across species revealed shared identity, as expected, but subsequent alignment of subtypes revealed varying patterns of conservation. A recent study of teleosts separated by 150 million years of evolution found that progenitor cell populations have higher conservation between species than differentiated cell populations^4^. Further, taxon-specific cell types (like the shell gland cells of oysters) were found to be the youngest transcriptomically of all the differentiated cell types in a scRNA-seq comparison within spiralia^37^. In support of these results, we found high conservation of cnidocyte progenitors across all cnidarians, representing ∼800 million years of evolution^17^ and, on average, expression of younger genes in cnidocytes. Comparison of differentiated cnidocytes, however, revealed varying degrees of alignment. We expected that the amount of time separating *A*. *poculata* and *N. vectensis* would have resulted in distinct transcriptional identities in homologous types of cnidocytes due to cellular systems drift, accumulated variation in gene expression that does not change cell function^4,50^. Surprisingly, we found extreme conservation of spirocyte gene expression and identified the deeply conserved transcription factor *FoxL2* as a critical regulator of spirocyte fate. Among nematocytes, the basitrichous isorhizas that are found in the embryonic ectoderm of both species were also highly conserved; in contrast, mastigophores showed almost no conservation of gene expression. In fact, mastigophores from *A. poculata* were found to be more similar to the stenoteles of *H. vulgaris* than the mastigophores of *N. vectensis*. Mastigophores are found only in the pharynx and mesenteries of *N. vectensis*^16,25^ where they are used for prey immobilization but are found in the outer ectoderm in the embryos of *A. poculata*. If mastigophores in *A. poculata* and *N. vectensis* are used for different functions they may have experienced different selection pressures, providing a possible explanation for the lower conservation in gene expression between these two species. Alternatively, it is possible that the mastigophore morphology has evolved multiple times in hexacorals, reflecting a recent analysis that found mastigophore morphology to be convergent in anthozoans and medusozoans^8^. Future studies connecting gene expression to the fine control of cnidocyte morphology will be necessary to evaluate hypotheses about convergence in cnidocyte identity.

Our understanding of animal developmental is limited by our reliance on a few, well-established model systems in which gene manipulation is possible. To better understand how genetic regulation of homologous traits diverges over evolutionary time, it is essential to expand access to a wider variety of model organisms^51^. The northern star coral, *Astrangia poculata,* is a new model organism that is suitable for studies across a breadth of scientific fields^52^. Our work, in combination with recent publication of the complete genome^53^ and development of methods for manipulating gene function^21^, elevate *Astrangia poculata* as a critical model for investigating the development and evolution of cell identity in early-diverging animals. The comparison of cnidocytes between *A. poculata* and *N. vectensis* points to different evolutionary processes acting on different yet related cell types. Nematocytes are used to pierce and envenomate prey, and the different stinging apparatus morphologies have long been thought to be associated with specialization for different prey types^14,15^. This suggests that nematocytes may be evolving under diversifying selection enabling different species to exploit novel prey resources. Alternatively, small changes that impact the size or placement of spines along the eversible tubule may create variability without negatively impacting the function of a nematocyte; therefore, nematocytes may be diverging due to purely neutral processes. Unlike the diversity of modern nematocytes, spirocytes have maintained a highly conserved morphology, despite being found throughout hexacorals^18^. An entire nomenclature system has been developed to capture the diversity of nematocyte types^45^, but spirocytes can only be divided into two types that differ in the shape of their capsule^16^. The extreme conservation of transcriptional and morphological identity suggests spirocytes may be evolving under purifying selection. This could suggest that very few changes to the genetic program driving spirocyte development can have large negative impacts on the function of this cell type. This is supported by our observation that gracile spirocytes have an older transcriptional age than nematocytes, suggesting that the incorporation of novel genes into the spirocyte gene regulatory network may negatively impact cell function. With access to this new anthozoan model for cellular development, future studies could test the relationship between gene expression and the ability of spirocyte tubules to discharge from the capsule and stick to prey to evaluate this hypothesis. Due to this observed evolutionary stasis of both morphology and gene expression, we propose that spirocytes might be considered a ‘living fossil’ cell type.

## Acknowledgements

The authors are grateful for support from the Marine Biological Laboratory’s Whitman Fellowship program. We thank the past and current members of the Babonis, Reed, Womack, Pennell, Sharp, and Roberson laboratories for their helpful discussion of the results and feedback on the manuscript. We also thank Bill Grossman, Dr. Sean Grace, Kerry Bresnahan, Zoe Dellaert, and Dr. Jill Ashey for help in diving and collecting corals.

## Author’s Contributions

Conceptualization: SA, LSB

Methodology: SA, RMB, JFW, KS, LMR, LSB

Investigation: SA, RMB, JFW, LSB

Resources: JFW, KS, LMR, LSB

Funding acquisition: SA, LSB

Project administration: SA, LSB

Supervision: LSB

Writing – original draft: SA, LSB

Writing – review & editing: SA, RMB, JFW, KS, LMR, LSB

## Data, code, and materials availability

Code and R objects are available at: https://github.com/Babonis-Lab/Arnold-et-al-2026. scRNA-sequencing data is publicly hosted on the UCSC Cell Browser at: https://astrangia-dev-atlas.cells.ucsc.edu. All other data are available in the main text or the supplementary materials.

## Materials and Methods

### Animal care and maintenance

*Astrangia poculata* colonies were collected via SCUBA from Fort Wetherill State Park (depth ∼5-10 m) in Jamestown, RI. Colonies were collected under a Rhode Island Department of Environmental Management collector’s permit granted to Roger Williams University (permits 825 and 943) and transferred to the Marine Biological Laboratory in Woods Hole, MA, where they were kept in sea tables with natural seawater. Spawning was induced by rapidly raising the water temperature by ∼8°C in benchtop containers, as previously described^21^. After spawning, colonies were returned to the flowthrough system at 19-22°C and maintained in a 12:12 light cycle until the next spawn. Colonies were fed mysid shrimp (from brineshrimpdirect.com) the day prior to spawning.

*Nematostella vectensis* was kept in the dark at 16°C in 1/3X (12ppt) artificial seawater made with Tropic Marin Pro Reef Salt Mix (Tropic-marin.com #232008) diluted in deionized (DI) water. Animals were spawned as previously described^24,54^. Embryos were de-jellied as previously described^55^. Embryos were raised at 24°C.

### Tissue collection

*A. poculata* embryos were raised at room temperature (19-22°C) and fixed in 4% paraformaldehyde (PFA) diluted in seawater by directly adding 125ul of 32% PFA to 875ul of embryos in seawater, as previously described^21^. Embryos were fixed for 1 hour at room temperature or overnight at 4°C and then rinsed 3 times for 10 minutes each in 1X Phosphate Buffered Saline containing 0.1% Tween 20 (PTw). Embryos fixed for immunohistochemistry (IHC) were stored in PTw at 4°C. Embryos fixed for in situ hybridization (ISH) were then washed 1 time in sterile water, 2 times in 100% methanol, and then stored at -20°C in clean 100% methanol.

*N. vectensis* embryos were immobilized in 7% MgCl_2_ made in DI water and diluted equal parts with 1/3X seawater for up to 10 minutes prior to fixation. They were then fixed in 4% PFA with 0.25% glutaraldehyde for 1 minute at RT and further fixed in 4% PFA without glutaraldehyde for 1 hour at 4°C. Embryos were then rinsed 3 times for 5 minutes each in PTw, and samples fixed for ISH and IHC were washed and stored as described for *A. poculata*.

### Immunohistochemistry

Immunohistochemistry was performed in *A. poculata* as previously described in *N. vectensis*^56^. A commercial α-acetylated tubulin antibody (Sigma T6743) was used at a concentration of 1/200 in *A. poculata*. The NvGLWamide antibody^57^ was used at a concentration of 1/200 in both *A. poculata* and *N. vectensis.* The FMRFamide antibody (BMA Biomedicals T-4322) labels RFamide neurons in *N.vectensis*^58^ and was used in *A. poculata* at a concentration of 1/100. All antibodies were diluted in 5% normal goat serum (Sigma G9023-10ML) made in 1X phosphate buffered saline containing 0.2% Triton-X and 1% BSA (PBT). Samples were incubated in secondary antibody (Invitrogen A11004, Invitrogen A21245) diluted 1/500 in 1X phosphate buffered saline containing 0.2% Triton-X (PTx). F-actin was stained with phalloidin (Invitrogen A12379) for 30 minutes at room temperature at a concentration of 1/100 in PTx. Nuclei were counterstained with DAPI (Sigma 45-D9542) at a concentration of 1.43μM diluted in 1X Tris Buffer for 30 minutes at room temperature. Samples were imaged on a Zeiss LSM 900 confocal microscope and adjusted for brightness and contrast in Fiji^59^.

### Cloning, probe synthesis, and in situ hybridization

RNA extraction for *N. vectensis* was performed as previously described^60^. *A. poculata* embryos were homogenized by vortexing and preserved for RNA extraction in TRI reagent (T9424, Sigma, USA) at -80°C. RNA was extracted using the Zymo RNA Miniprep Kit (#76020-642) following the manufacturer’s protocol. cDNA synthesis was performed using the Takara Advantage RT-for-PCR Kit (639506) following the manufacturer’s protocol. Probe synthesis and mRNA in situ hybridization for both *A. poculata* and *N. vectensis* were performed as previously described^21,60^. Primers used to generate probes can be found in **Table S1**. At the time of use, mRNA probes were diluted in hybridization buffer to a working concentration of 1ng/uL for *N. vectensis* and 5ng/uL or 1ng/uL (probe dependent) for *A. poculata*. ISH was performed in *A. poculata* following the protocol as described in Warner et al. (2025)^21^ with minor modifications. Baskets were made from 2mL screw cap tubes cut in half with 30 μm mesh secured in the lid and were used to move embryos between washes in a sterile 24 well plate. Proteinase-K digestion was done for only 1-2 minutes, and an increased probe concentration (5ng/uL) was used. Samples were imaged on a Nikon Eclipse E800 microscope, scale bars were added in Fiji^59^, and brightness was adjusted in Adobe Photoshop 2025.

### Squash preparations and cnidocyte counting

*A. poculata* embryos previously fixed and stored at 4°C in PTw were re-fixed in 4% PFA overnight at 4°C and then rinsed in PTw 3 times for 5 minutes each. Individual embryos were mounted on glass slides in a minimal amount of 80% glycerol diluted in 1X PBS. Coverslips were then pressed down on the embryo and smeared to break apart all tissue except for the remaining hardened cnidocyte capsules. Slides were sealed with nail polish to create a permanent mount. The edges of the dissociated tissue for each sample were identified, and images were taken across the entirety of the sample using the tiling function taking on a Zeiss LSM 900 confocal microscope. Tiled images were then stitched into one image per embryo using Zen 3.8 software. All cnidocytes in each image were counted manually using the Cell Counter function in Fiji as well as measured for length and width using the measure function in Fiji^59^. Raw cnidocyte counts can be found in **Table S2** and cnidocyte measurements can be found in **Table S3**.

### Minicollagen tree

Minicollagen sequences were identified from the *A. poculata* predicted proteome using a reciprocal best blast strategy. Amino acid sequences for all five minicollagens from *N. vectensis* (mcol1, mcol3, mcol4, mcol5, and mcol6) were downloaded from SIMRbase and searched against the *A. poculata* translated transcriptome using blastp. Top hits were reciprocally searched against the proteome of *N. vectensis* (NV2 version, downloaded from SIMRbase) to find one-to-one orthologs. The resulting sequences were determined to be minicollagens only if they contained one or more central glycine-X-Y domain(s) flanked by polyproline repeats and cysteine rich domains^40^. Verified minicollagen sequences were then added to an existing alignment of minicollagens from across cnidarians^61^ using mafft^62^. To find the best tree, we performed three maximum likelihood analyses in parallel using RAxML v.8^63^ and IQ-TREE v.3^64^: RAxML with 25 random starting trees, RAxML with 25 maximum parsimony starting trees, and IQ-TREE with default parameters. All trees were generated using a PMB substitution matrix, identified using the model finder feature in IQ-TREE3. Outputs were evaluated by their maximum likelihood scores and the IQTREE was chosen. Ultrafast bootstraps were added using the -bb 1000 tag. The final tree was produced using the same protocol after the alignment was trimmed to include only *N. vectensis* and *A. poculata*. Tips were aligned using FigTree v.1.4.4^65^.

### Manipulation of gene expression

FoxL2 knockdown in *N. vectensis* was conducted using short hairpin RNAs (shRNAs). shRNAs were synthesized following the protocol detailed in Karabulut et al. 2019^47^. shRNA primer sequences can be found in **Table S1**. Gene-specific and scrambled control shRNAs were injected into zygotes at a concentration of 800ng/uL diluted in nuclease-free water (Fisher AM9937) with dextran (Fisher D34679) at a concentration of 1mg/mL^21^. After injection, embryos were raised at 24°C.

### Spirocyte labeling

Spirocytes in *N. vectensis* were labeled with a fluorescein tyramide used previously for fluorescent in situ hybridization in this species^44^. Embryos were permeabilized with 5 washes (10 minutes each) in PBT and then incubated in fluorescein tyramide diluted 1/500 in PTw for 1 hour at room temperature in the dark. Embryos were then washed extensively with PTw and nuclei were labeled with 1.43μM DAPI as described above. Samples were imaged on a Zeiss LSM 900 confocal microscope and adjusted for brightness and contrast in Fiji^59^. The number of spirocytes in each embryo was counted using the Cell Counter plugin in Fiji^59^ and the bar graph was generated using the ggplot2 (3.5.2) package^66^ in R.

### GLWamide-expressing neuron quantification

Samples were imaged on a Zeiss LSM 900 confocal microscope and adjusted for brightness and contrast in Fiji^59^. The number of GLWamide+ neurons in each embryo was counted on max projected images using the Cell Counter plugin in Fiji^59^ and the bar graph was generated using the ggplot2 (3.5.2) package^66^ in R. Statistical significance was determined by running a Wilcox rank sum test on the data in R. Raw data can be found in **Table S4**.

### Embryo dissociations and scRNA-sequencing

Dissociation and fixation of *A. poculata* embryos for scRNA-seq were carried out following a protocol adapted from Massri et al. 2021^67^. Embryos were collected every 12 hours after fertilization up to 96 hours post fertilization(hpf). Approximately 800 embryos were collected for the 12hpf sample and approximately 450-600 embryos were collected for each subsequent sample. Embryos were transferred to a glass 9-spot plate chilled on ice and washed 2 times each in 1mL of 2% N-acetyl-L-cysteine (NAC) (45-A7250, Sigma, USA) diluted in calcium/magnesium-free artificial seawater (CMF-ASW: 461.85 mM NaCl, 10.73 mM KCl, 7.04 mM Na_2_SO_4,_ 2.14 mM NaHCO_3_, pH: 8.2). Embryos were then dissociated in 100uL of 2% NAC in CMF-ASW via mechanical agitation by pipetting. Once dissociated, 900uL of chilled 100% methanol was added dropwise to the spot containing dissociated embryos in 2% NAC in CMF-ASW. The dissociated cell suspension was then transferred to a 1.5ml microcentrifuge tube coated in 1% BSA (diluted in sterile DI water) and stored at -20°C for less than one month prior to processing. In preparation for sequencing, cell suspensions were centrifuged at 1500g for 10 min at 4°C, the supernatant was removed, and then cells were rehydrated in a solution of 3x SSC (diluted in sterile DI water from 20x SSC, Sigma SRE0068) containing 1% BSA. Dissociated cells were then filtered through a SP Bel-Art Flowmi 40 um filter (VWR 10032-802) to remove clumps. Filtered cells were then used immediately for library preparation. Library preparation and sequencing were performed by the Biotechnology Resource Center (BRC) Genomics Facility (RRID:SCR_021727) at the Cornell Institute of Biotechnology. Libraries for each timepoint were prepared using the 10x Genomics Chromium GEM-X 3’ Gene Expression v4 kit (PN-1000691) and sequenced using a NovaSeqX 25B lane. Mean reads per cell for each timepoint can be found in **Table S5**.

### Single-cell RNAseq analysis

#### Whole developmental atlas

10x Genomics Cell Ranger v8.0.0^68^ was used to make a reference genome with the ‘mkref’ command using the *A. poculata* genome generated by Stankiewicz et al. (2025)^53^. The previously published genome annotations were extended to include 3’UTR regions using GUSHR to improve genome mapping of reads^69–74^. Reads were then mapped to the reference genome using the ‘count’ command with default parameters. The 24hpf timepoint was forced to have a recovery of 7000 cells as there was some cell clumping in this sample causing default parameters to recover too few cells. The resulting h5 output files were read into R and a Seurat object was made for each timepoint using Seurat 5.4.0^75^. The objects were then merged and the following was done using Seurat 5.4.0^75^. Low-quality cells were filtered out using nFeature_RNA > 20 & nFeature_RNA < 3500 based on the generated violin plots. Following filtering, the data were normalized and scaled, and variable features across cells were found. We performed principal component analysis (PCA) on the scaled data and visualized the results using an elbow plot. We chose to use the first 15 PCs as they had the highest standard deviation to construct a k-nearest neighbor (KNN) graph. Clustering was performed with the default resolution of 0.5 and visualized using Uniform Manifold Approximation and Projection (UMAP). Genes that were upregulated in each cluster were identified using the ‘FindAllMarkers’ function and are available in **Table S6**. Clusters were annotated by projecting known cell type genes from *N*. *vectensis* onto the UMAP and validated via in situ hybridization. Targets for validation by in situ hybridization were chosen from the set of genes for each cluster with the highest avg_log2FC that were also specifically expressed in only the cluster of interest. The population “progen.1” is made up of two clusters and the “sec.progen.2” population is made up of four clusters. We chose to group clusters determined by Seurat because they did not strongly express any genes uniquely that we could use to justify calling them different cell populations. Progenitor clusters were annotated after projecting markers onto the developmental trajectory URD tree.

#### Cnidocyte subset

Cnidocyte clusters were extracted from the whole developmental atlas and processed using the same Seurat pipeline described above. Clustering was performed with a resolution of 0.4 as this best recapitulated the number of cnidocytes we determined empirically using morphological analysis. Clusters were annotated by projecting known cnidocyte genes onto the UMAP and comparing the timepoints that make up each cluster to the known time of each cnidocyte type development. Clusters were validated with in situ hybridization of marker genes chosen as described above. Genes that were upregulated in each cluster were identified using the ‘FindAllMarkers’ function and are available in **Table S7**.

#### Neuron + gland cell subset

Neuron + gland cell progenitors and differentiated clusters were extracted from the whole developmental atlas and processed using the same Seurat pipeline described above. Clustering was performed using the default resolution of 0.6. Clusters were annotated by projecting marker genes from the whole developmental atlas onto the UMAP. Clusters that expressed Ap.2.3087 (neural marker) were annotated as N1.X and those that expressed the neural marker *AshA* (Layden et al. 2012) were annotated as N.X. Cluster NGD.progen expressed Apoc2.1-6.919 (neurogland.progen marker). The top markers of clusters annotated NGD.early.X were projected onto the developmental URD tree and found to be expressed highly and early in the branch leading to neurons and gland cells. Clusters that expressed the markers for sec.progen.1 and sec.progen.2 were annotated as such. The cluster S.early was annotated as such because the top expressed genes in this cluster are expressed in the branch leading to secretory cells populations gland.1, gland.mouth, secretory.2, and secretory.3. Clusters that expressed the markers for gland.1, secretory.1, secretory.2, and secretory.3 were annotated as G1, S1, S2, and S3, respectively. Clusters that expressed CHIT1-like-13 (gland.mouth marker) were annotated as GM.X and clusters that expressed Apoc2.1-9.505 (mucous.gland marker) were annotated as M.X. The cluster that expressed markers of cell division was annotated as pSCs. Remaining clusters were annotated as NGD.X. Four clusters were grouped to make the sec.progen.1 population. Genes that were upregulated in each cluster were identified using the ‘FindAllMarkers’ function and are available in **Table S8**.

### URD developmental trajectory analysis

Differentiation trajectory analysis was performed with the R package URD following the authors’ recommendations^32^. To generate the developmental URD tree, the blastula cluster was set as the root of the tree, and the 10 differentiated cell type clusters were set as the tips of the tree. The mature cnidocyte cluster was not included as all cnidocyte types converge on a single homogeneous transcriptional identity^25,27^. Many parameters were tried and a sigma of 30 and knn of 100 were ultimately determined as optimal by examining diffusion maps. We were only able to recover one lineage of spirocytes in the developmental URD tree due to the small number of gracile spirocytes. In order to gain better resolution of cnidocyte development, we used the cnidocyte subset data to construct a lineage trajectory tree. The early progenitor cluster is made up of only 24hpf cells and expresses the cnidocyte progenitor marker *CnZnf1*^43^; therefore, it was set as the root of the tree and the four cnidocyte clusters were set as the tips of the tree. The mature cnidocyte cluster was not included as described above. NULL sigma and knn of 50 were determined as optimal by examining diffusion maps. Results of differential expression analysis for the spirocyte branch can be found in **Table S9**.

### SAMap analysis

The *A. poculata* transcriptome^53^ and the *N. vectensis* NV2 transcriptome (available at: https://simrbase.stowers.org/starletseaanemone) were searched against one another using blastn to identify one-to-one orthologs. Cell count matrices and metadata files containing Seurat clustering information were extracted from our *A. poculata* cnidocyte and neuron + gland cell subsets, from available *N. vectensis* cnidocyte, neuron + gland cell, and mucous cell subsets^25,26^, and from the available *H. vulgaris* interstitial subset^27^. *Xenia sp.* cnidocyte clusters were generated by subclustering the scRNA-seq dataset including homeostatic and regeneration data generated by Hu et al. (2020)^29^. Clusters were annotated based on expression of cnidocyte progenitor and mature cnidocyte markers from *A. poculata* and *N. vectensis*. Barcode and feature files were extracted for each species and used to generate hierarchical data format files (.h5ad) using the AnnData (0.11.4) Python package^76^. SAMap v.1.0.16^3^ was run following the program’s documentation using the BLAST outputs and .h5ad files to integrate the datasets together. Cnidocyte and neuron + gland cell subset .h5ad files for *A. poculata* and *N. vectensis* were concatenated and run through SAMap so that alignment scores could be accurately compared across cnidocytes, neurons, and gland cells. For broad cell type comparisons, cnidocyte and neuron + gland cell subclusters were collapsed into broader categories (e.g. basitrichous isorhizas and mastigophores were collapsed into the nematocyte category) and aligned using SAMap. Cell type mapping scores were calculated using the SAMap get_mapping_scores function which were used to generate a heatmap using the R pheatmap (1.0.13) package^77^ or ComplexHeatmap (2.27.1)^78,79^ or Sankey plots using the R networkD3 (0.4.1) package^80^. Alignment scores for broad cell type comparisons between *A. poculata* and *N. vectensis,* neuroglandular subpopulation comparisons between *A. poculata* and *N. vectensis*, cnidocyte comparisons between *A. poculata, N. vectensis,* and *Xenia sp.*, and neuroglandular comparisons between *A. poculata, N. vectensis,* and *H. vulgaris* can be found in **Table S10, Table S11, Table S12, and Table S13,** respectively. Gene pairs enriched in pairs of cell types were identified using the SAMap GenePairFinder function. Enriched gene pairs for broad cell type comparisons between *A. poculata* and *N. vectensis,* neuroglandular subpopulation comparisons between *A. poculata* and *N. vectensis*, cnidocyte comparisons between *A. poculata, N. vectensis,* and *Xenia sp.*, and neuroglandular comparisons between *A. poculata, N. vectensis,* and *H. vulgaris* can be found in **Table S14, Table S15, Table S16, and Table S17,** respectively. The number of genes expressed in each cluster as well as the number of cells in each cluster were extracted from the neurogland and cnidocyte Seurat objects for both *A. poculata* and *N*. *vectensis.* Plot was made using ggplot2 (3.5.2) package^66^ in R. Raw data is available in **Table S18**. Functional enrichment analysis was performed on each cluster using the KOG (Eukaryotic Orthologous Groups) terms generated from running each species’ transcriptome through EggNOG-mapper (2.1.13)^81^. Tables generated by EggNOG-mapper for *A. poculata* and *N. vectensis* can be found in **Table S19** and **Table S20**, respectively.

### Phylostratigraphy analysis

Phylostratigraphy analyses were adapted from Piovani et al., 2023^37^. The *A. poculata* transcriptome^53^ and the *N. vectensis* NV2 transcriptome (available at: https://simrbase.stowers.org/starletseaanemone) were used as input into the phylostratigraphy program GenEra^38^ following the provided documentation to estimate the age of each gene. With the R package myTAI^39^, we calculated the transcriptome age indices (TAI) of each cell type using the gene phylostratum outputs for both species from GenEra and the log-transformed gene average expression of each cluster that was calculated using the Seurat function ‘AverageExpression’. Plot were generated using ggplot2 (3.5.2) package^66^ in R.

## Table Captions

**Table S1:** List of primers used to make riboprobes for in situ hybridization and shRNAs.

**Table S2:** Raw counts of the different cnidocyte types at each timepoints in *A. poculata* embryos.

**Table S3:** Length and width measurements of each cnidocyte type in *A. poculata* embryos.

**Table S4:** Quantification of GLWamide+ neurons in control and FoxL2 KD embryos

**Table S5:** Number of raw reads, estimated number of cells, and mean reads per cell generated from Cell Ranger count on whole developmental atlas.

**Table S6:** Marker genes generated using FindMarkerGenes in Seurat for whole developmental cell atlas

**Table S7:** Marker genes generated using FindMarkerGenes in Seurat for cnidocyte scRNA-seq subset

**Table S8:** Marker genes generated using FindMarkerGenes in Seurat for neurogland scRNA-seq subset

**Table S9:** Differentially expressed genes in the branch leading to spirocytes in the *A. poculata* cnidocyte URD tree.

**Table S10:** Alignment scores generated in SAMap comparing broad neuroglandular cell types between *A. poculata* and *N. vectensis*.

**Table S11:** Alignment scores generated in SAMap when comparing neuroglandular subpopulations between *A. poculata* and *N. vectensis*.

**Table S12:** Alignment scores generated in SAMap when comparing cnidocytes between *A. poculata*, *N. vectensis*, and *Xenia sp*.

**Table S13:** Alignment scores generated in SAMap when comparing neuroglandular cells between *A. poculata*, *N. vectensis*, and *Hydra vulgaris*.

**Table S14:** Gene pairs for cell type cluster pairs generated in SAMap when comparing broad neuroglandular cell types between *A. poculata* and *N. vectensis*.

**Table S15:** Gene pairs for cell type cluster pairs generated in SAMap when comparing neuroglandular subpopulations between *A. poculata* and *N. vectensis*.

**Table S16:** Gene pairs for cell type cluster pairs generated in SAMap when comparing cnidocytes between *A. poculata*, *N. vectensis*, and *Xenia sp*.

**Table S17:** Gene pairs for cell type cluster pairs generated in SAMap when comparing neuroglandular cells between *A. poculata*, *N. vectensis*, and *Hydra vulgaris*.

**Table S18:** The total number of genes expressed and number of cells in each neuroglandular cluster in *A. poculata* and *N. vectensis*.

**Table S19:** Output generated from running the *A. poculata* transcriptome through EGGNOG that was then used for functional enrichment analysis in SAMap.

**Table S20:** Output generated from running the *N. vectensis* transcriptome through EGGNOG that was then used for functional enrichment analysis in SAMap.

---

**Supplemental Figure S1:**
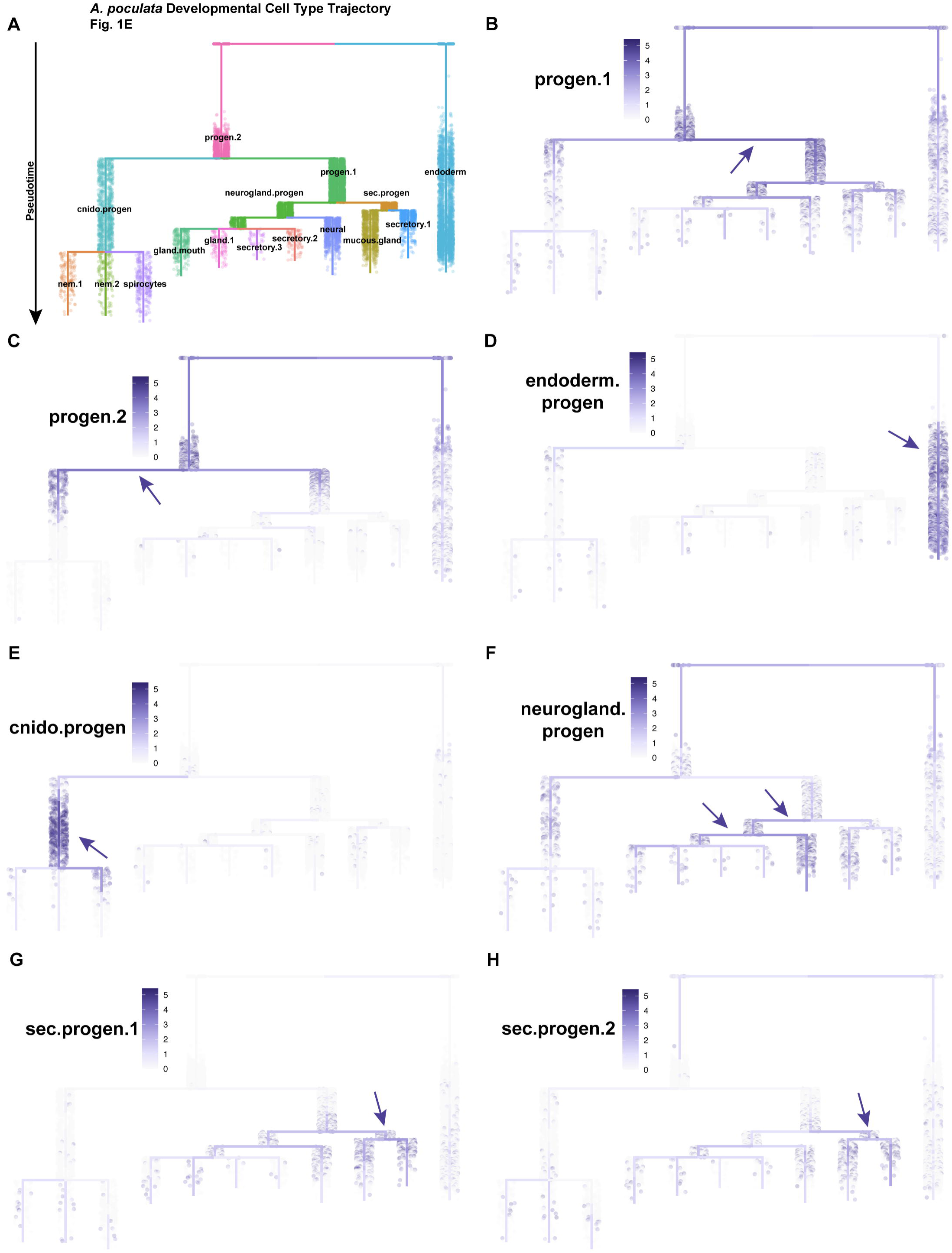
Visualization of progenitor cell trajectories throughout development in *A. poculata*. **A)** Developmental cell trajectory generated in URD shown in Figure 1E. **B-H)** Projection of progenitor cell markers from Figure 1G-J on *A. poculata* developmental trajectory tree. Arrows point to highest expression.

**Supplemental Figure S2:**
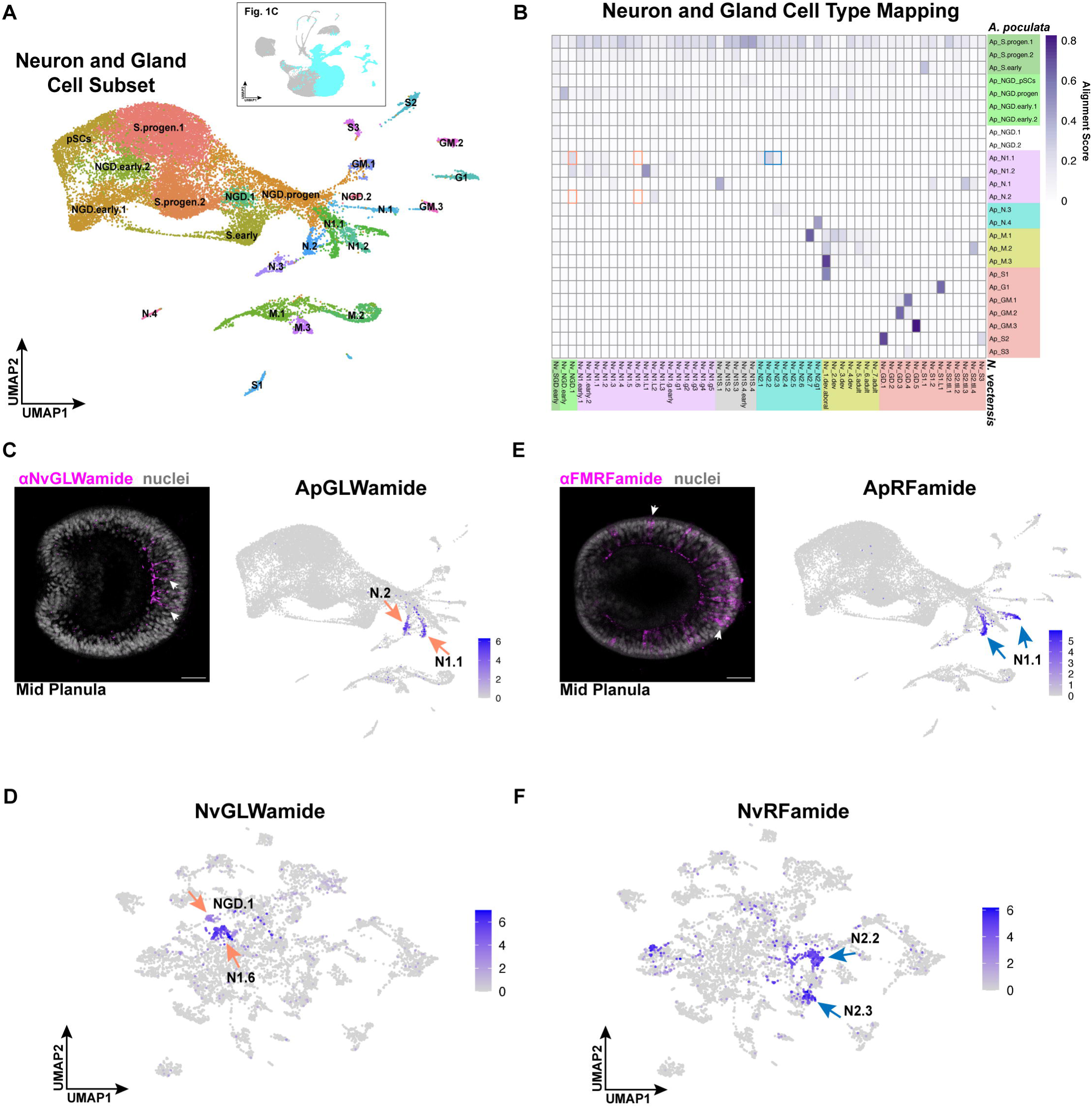
Aligning *A. poculata* and *N. vectensis* neurons and gland cell subsets identifies few cell types with high conservation. **A)** Neuron and gland cell subset UMAP of *A. poculata.* NGD = neuroglandular, N = neural, M = mucous, S = secretory, and GM = gland mouth. **B)** Heat map of alignment scores generated in SAMap. Colored boxes correspond to cell categories in Figure 2B. GLWamide-expressing neurons and RFamide-expression neurons are highlighted in orange and blue, respectively. **C)** Expression of *ApGLWamide* precursor projected onto neuron and gland cell subset UMAP. GLWamide+ neurons are labeled in magenta in *A. poculata* mid planula embryos using a *N. vectensis* GLWamide antibody (Nakanishi and Martindale 2018). Nuclei are labeled with DAPI in grey. **D)** Expression of *NvGLWamide* precursor projected onto neuron and gland cell subset UMAP^25,26^. **E)** Expression of *ApRFamide* precursor projected onto neuron and gland cell subset UMAP. Expression of *ApRFamide* RFamide+ neurons are labeled in magenta in *A. poculata* mid planula embryos using a commercial FMRFamide antibody. Images are max projections of a few slices at the midpoint of the embryo. **F)** Expression of *NvRFamide* precursor projected onto neuron and gland cell subset UMAP 25,26.

**Supplemental Figure S3:**
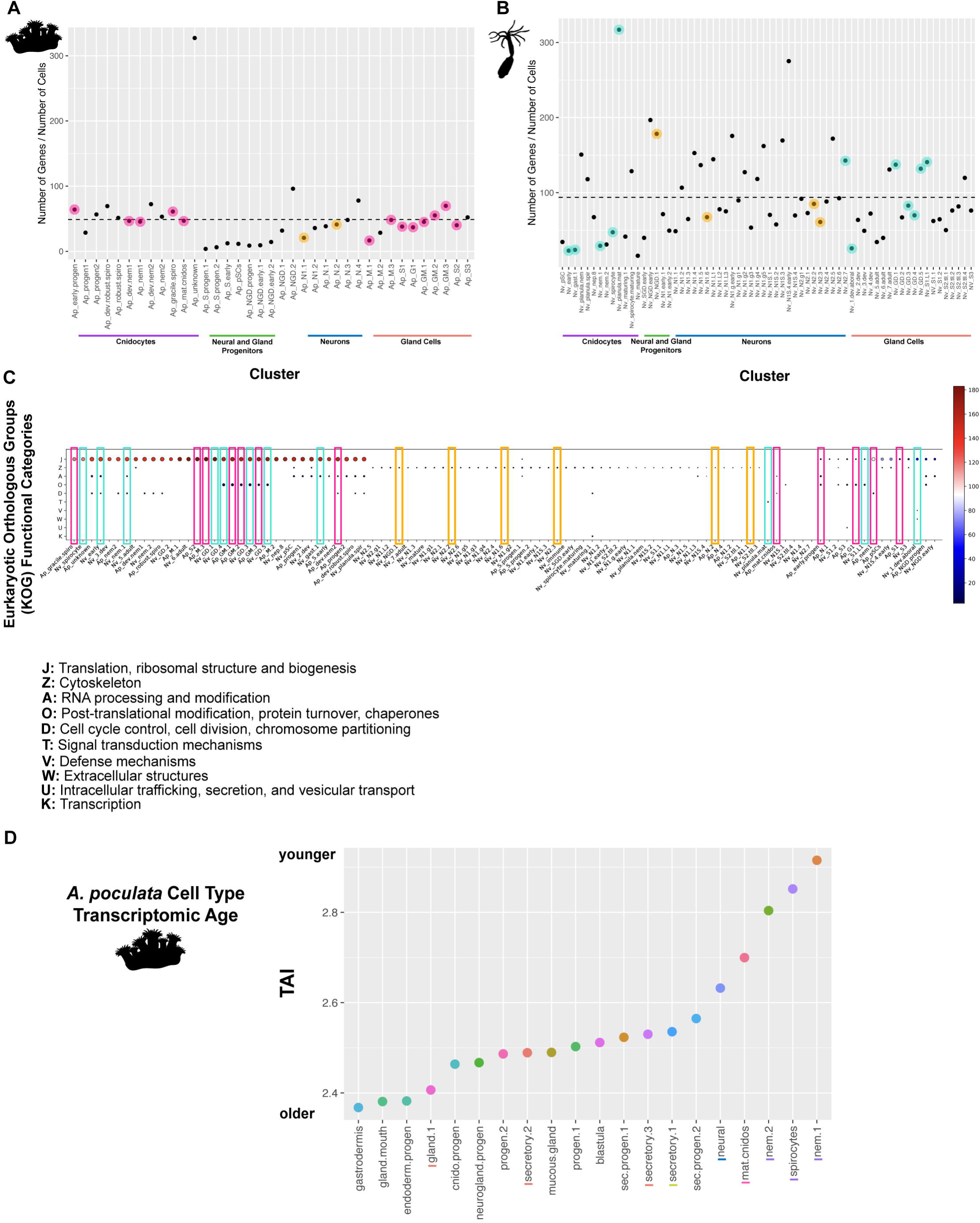
Assessment of identity-driven genes in neuroglandular cell clusters. **A-B)** Because more genes may be detected in clusters with more cells, we normalized the number of genes expressed to the number of cells comprising each cluster in *A. poculata* (A) and *N. vectensis* (B). Black dashed line indicates mean normalized number of genes per cluster. Clusters that aligned highly between species are highlighted in pink (*A. poculata*) and teal (*N. vectensis*). GLWamide+ and RFamide+ neural clusters are highlighted in orange. **E)** Functional categories of enriched genes in each cell type cluster. Clusters are highlighted as described in (A-B). Dot color represents degree of enrichment for each KOG term and dot size represents number of genes enriched for KOG term in each cluster. **D)** Analysis of gene age in each cell type in *A. poculata*. Y-axis shows transcription age index (TAI), with larger TAI values corresponding to younger gene ages. Dot color corresponds to cell type color in Fig. 1C. Lines underneath cell type name highlight those compared with *N. vectensis* in Fig. 2B and C, with colors corresponding to those highlighting cell types in Fig. 2B.

**Supplemental Figure S4:**
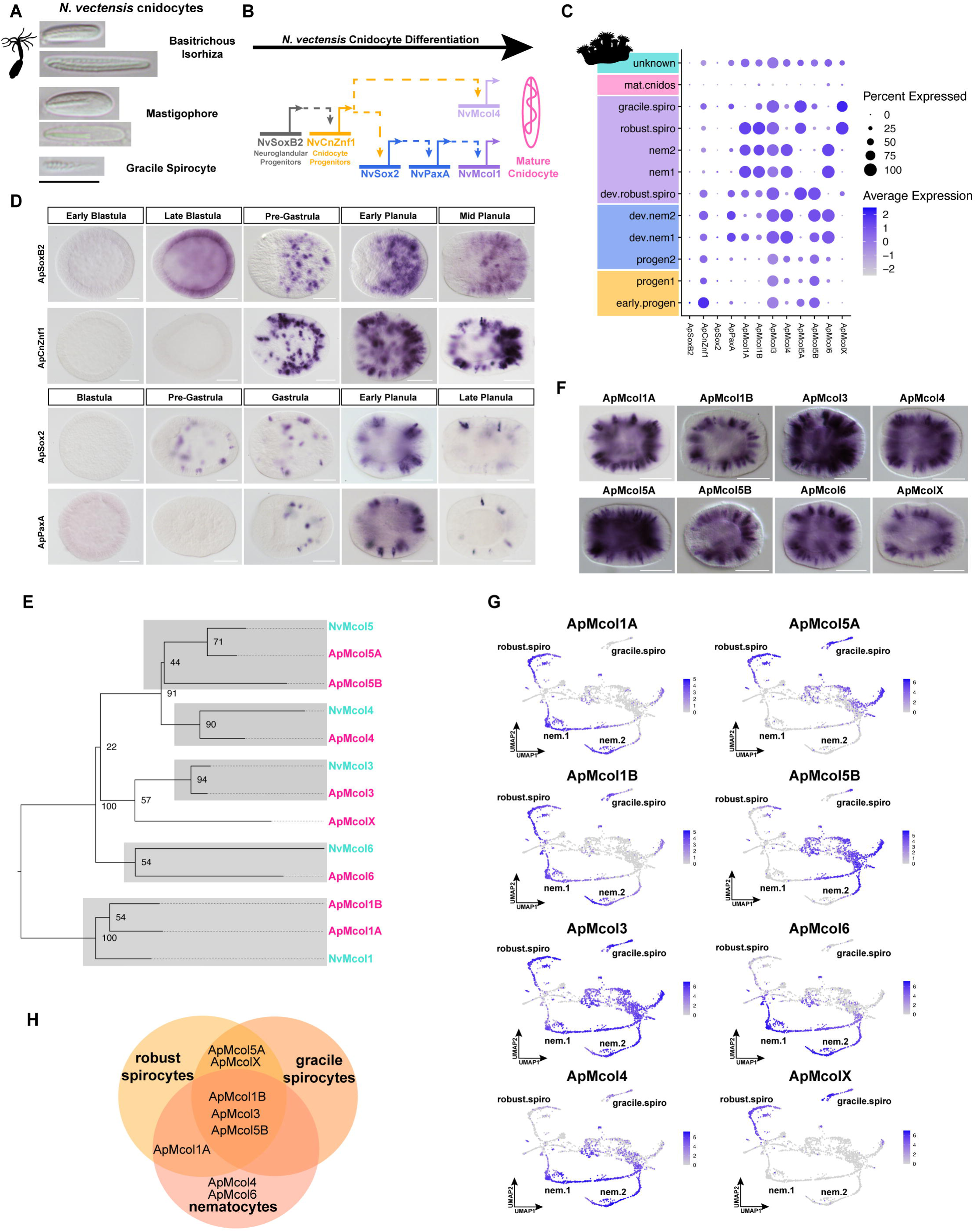
Comparison of *A. poculata* and *N. vectensis* cnidocytes with those of an octocoral (*Xenia sp.*) and a medusozoan (*Hydra vulgaris*) shows minimal alignment of cnidocytes. **A)** Heat map showing alignment scores generated in SAMap for all cnidocyte types in *A. poculata* (pink)*, N. vectensis* (teal), and *Xenia sp*. (blue). Colored boxes correspond to broad cnidocyte categories as in Figure 2C. *N. vectensis* and *A. poculata* comparisons are the same data shown in Figure 2C. **B)** Heat map showing alignment scores generated in SAMap for all cnidocyte types in *A. poculata* (pink)*, N. vectensis* (teal), and *H. vulgaris* (orange). **C)** Heat maps in A and B show cnidocyte alignments for animals with the relationships show in the tree. Progenitors and mature cnidocytes align across all species.

**Supplemental Figure S5:**
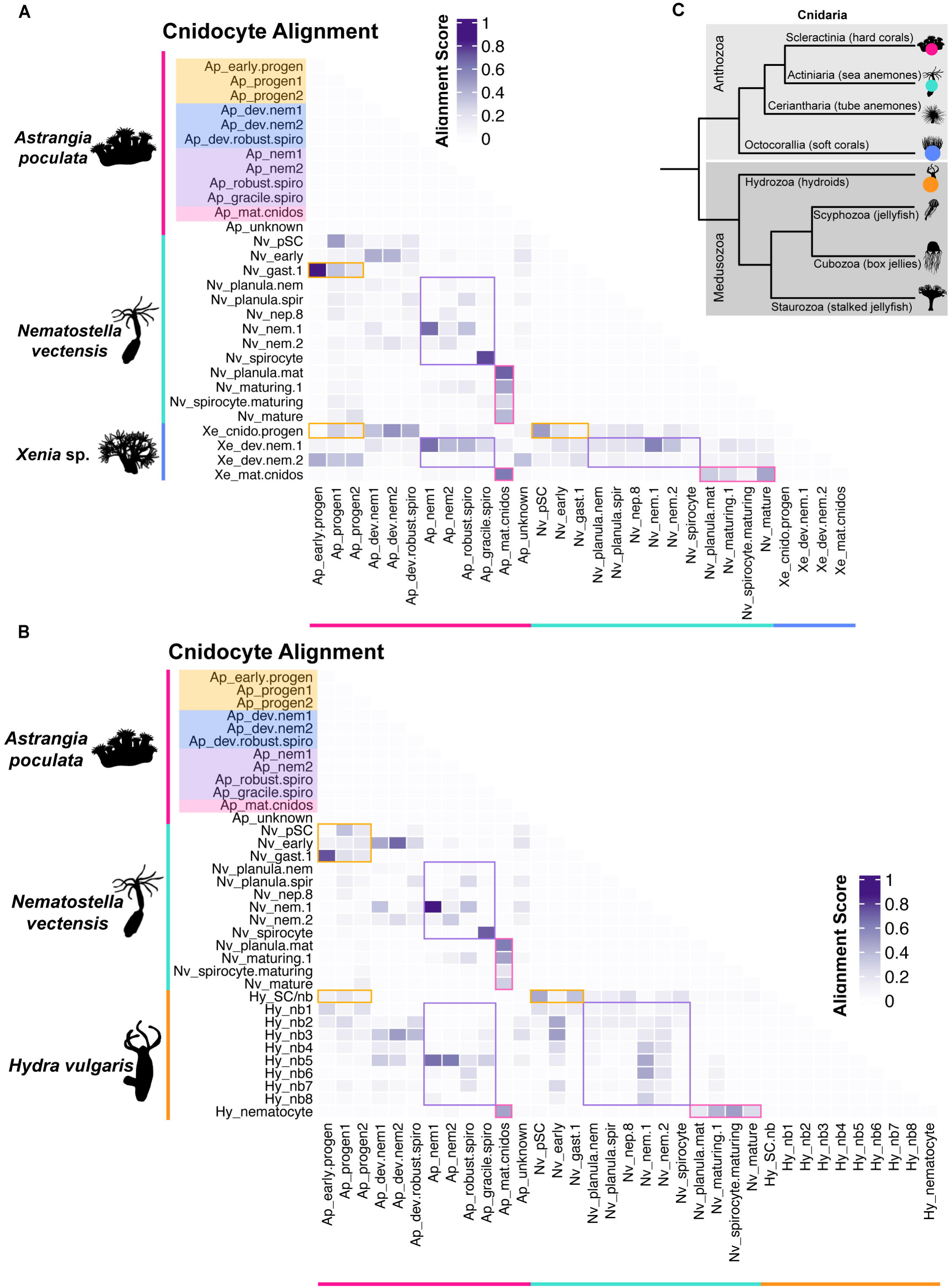
Knockdown of *NvFoxL2* results in complete loss of *NvCaluF* and small decrease in NvGLWamide. **A)** In situ hybridization of *ApFoxL2* in early planula embryos. *ApFoxL2* expression projected on scRNAseq UMAP (Fig 1C). Arrows indicate putative neurons identified by morphology. **B)** In situ hybridization of *NvFoxL2* in late planula embryos. *NvFoxL2* expression projected on scRNAseq UMAP^25,26^. Scale bars are 50μm for whole embryo images and 10μm for side panels. **C)** In situ hybridization of *NvFoxL2* in control embryos injected with scrambled shRNAs^47^ and embryos injected with *NvFoxL2* short hairpin RNAs (*NvFoxL2* KD). shRNAs are effective only until late planula stage, after which wildtype expression of *NvFoxL2* is restored. **D)** In situ hybridization of *NvCaluF* in control and *NvFoxL2* knockdown embryos. Primary polyp stages are shown in oral view with asterisk indicating the mouth. TB = tentacle bud, TT = tentacle tip. **E)** GLWamide labeled in magenta in control and *NvFoxL2* KD embryos. Nuclei labeled with DAPI in grey. Asterisks indicate mouth. **F)** There is a small but significant decrease in the number of GLWamide+ neurons in *NvFoxL2* KD embryos. Line in box plot shows mean and boxes around mean show standard deviation. GLWamide+ neurons in whole max projected embryos were counted. N = 12 control and 12 KD embryos. C-E All scale bars are 50μm.

**Figure S6.**
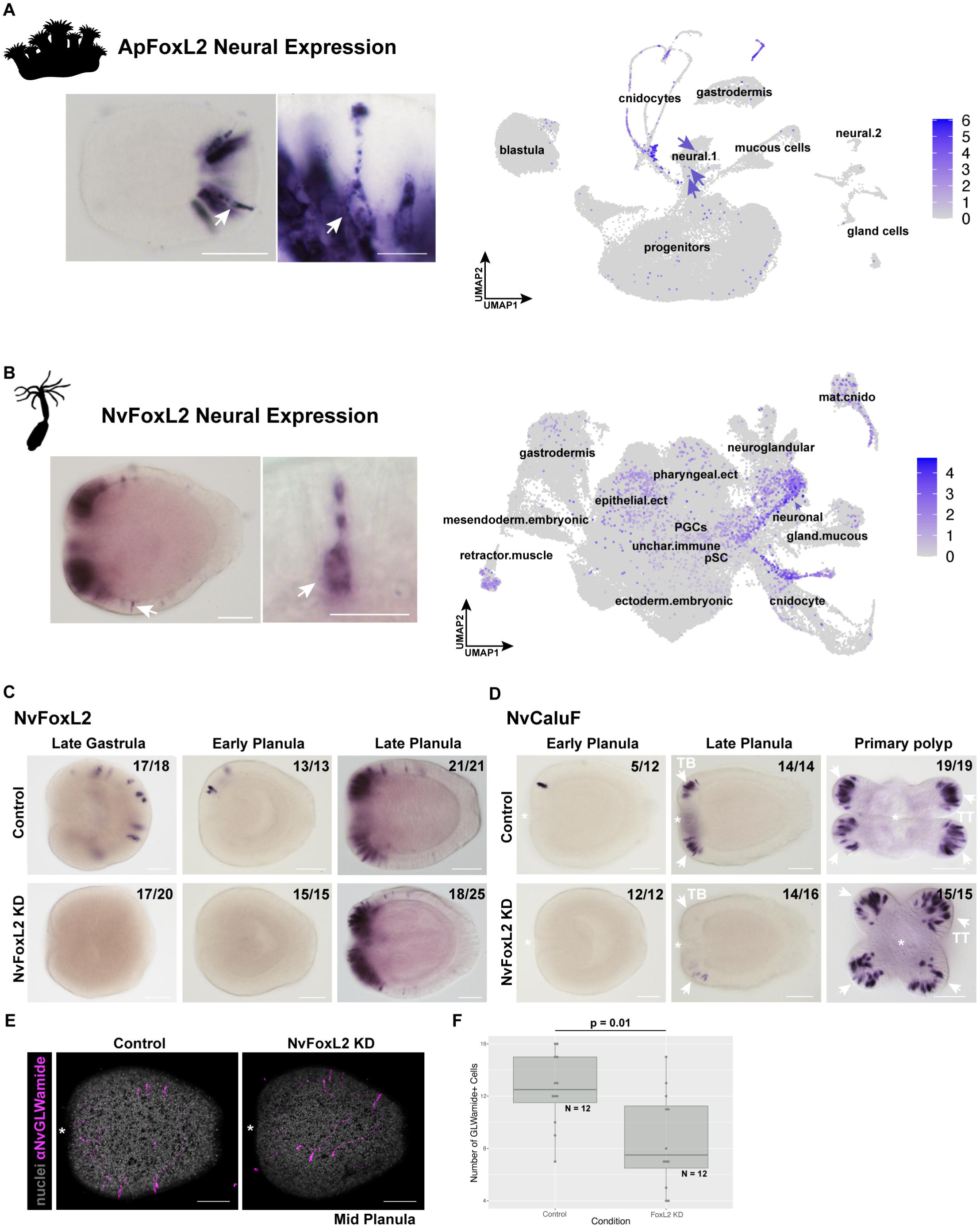

## References

1. Sebé-Pedrós, A. et al. Early metazoan cell type diversity and the evolution of multicellular gene regulation. *Nat*. Ecol. Evol. 2, 1176–1188 (2018).

2. Sebé-Pedrós, A. et al. Cnidarian Cell Type Diversity and Regulation Revealed by Whole-Organism Single-Cell RNA-Seq. Cell 173, 1520–1534.e20 (2018).

3. Tarashansky, A. J. et al. Mapping single-cell atlases throughout Metazoa unravels cell type evolution. eLife 10, e66747 (2021).

4. Shafer, M. E. R., Sawh, A. N. & Schier, A. F. Gene family evolution underlies cell-type diversification in the hypothalamus of teleosts. *Nat*. Ecol. Evol. 6, 63–76 (2022).

5. Hehmeyer, J., Harris, D., Lowe, C. & Marlow, H. Evolutionary principles underlying neuron subtype encoding and diversification in animals. 2026.04.13.718258 Preprint at 10.64898/2026.04.13.718258 (2026).

6. Tanay, A. & Sebé-Pedrós, A. Evolutionary cell type mapping with single-cell genomics. Trends Genet. 37, 919–932 (2021).

7. Dohrmann, M. & Wörheide, G. Dating early animal evolution using phylogenomic data. Sci. Rep. 7, 3599 (2017).

8. Sierra, N. C. & Gold, D. A. The evolution of cnidarian stinging cells supports a Precambrian radiation of animal predators. Evol. Dev. 26, e12469 (2024).

9. Mariscal, R. N. Cnidaria: Cnidae. in Biology of the Integument: Invertebrates (eds. Bereiter-Hahn, J., Matoltsy, A. G. & Richards, K. S.) 57–68 (Springer, Berlin, Heidelberg, 1984). doi:10.1007/978-3-642-51593-4_6.

10. Kass-Simon, G. & Scappaticci, Jr., A. A. The behavioral and developmental physiology of nematocysts. Can. J. Zool. 80, 1772–1794 (2002).

11. Beckmann, A. & Özbek, S. The Nematocyst: a molecular map of the Cnidarian stinging organelle. Int. J. Dev. Biol. 56, 577–582 (2012).

12. Holstein, T. & Tardent, P. An Ultrahigh-Speed Analysis of Exocytosis: Nematocyst Discharge. Science 223, 830–833 (1984).

13. Mapstone, G. M. Global Diversity and Review of Siphonophorae (Cnidaria: Hydrozoa). PLoS ONE 9, e87737 (2014).

14. Purcell, J. E. & Mills, C. E. The correlation between nematocyst types and diets in pelagic hydrozoa. in The Biology of Nematocysts 463–485 (1988).

15. Damian-Serrano, A., Haddock, S. H. D. & Dunn, C. W. The evolution of siphonophore tentilla for specialized prey capture in the open ocean. Proc. Natl. Acad. Sci. 118, e2005063118 (2021).

16. Babonis, L. S. et al. Single-cell atavism reveals an ancient mechanism of cell type diversification in a sea anemone. Nat. Commun. 14, 885 (2023).

17. McFadden, C. S. et al. Phylogenomics, Origin, and Diversification of Anthozoans (Phylum Cnidaria). Syst. Biol. 70, 635–647 (2021).

18. Fautin, D. G. Structural diversity, systematics, and evolution of cnidae. Toxicon 54, 1054–1064 (2009).

19. Kraus, Y. & Technau, U. Gastrulation in the sea anemone Nematostella vectensis occurs by invagination and immigration: an ultrastructural study. Dev. Genes Evol. 216, 119–132 (2006).

20. Magie, C. R., Daly, M. & Martindale, M. Q. Gastrulation in the cnidarian *Nematostella vectensis* occurs via invagination not ingression. Dev. Biol. 305, 483–497 (2007).

21. Warner, J. F. et al. Microinjection, gene knockdown, and CRISPR-mediated gene knock-in in the hard coral, Astrangia poculata. EvoDevo 16, 6 (2025).

22. Szmant-Froelich, A., Yevich, P. & Pilson, M. E. Q. Gametogenesis and Early Development of the Temperate Coral Astrangia danae (Anthozoa: Scleractinia). Biol. Bull. 158, 257–269 (1980).

23. Steinmetz, P. R. H., Aman, A., Kraus, J. E. M. & Technau, U. Gut-like ectodermal tissue in a sea anemone challenges germ layer homology. *Nat*. Ecol. Evol. 1, 1535–1542 (2017).

24. Hand, C. & Uhlinger, K. R. The Culture, Sexual and Asexual Reproduction, and Growth of the Sea Anemone Nematostella vectensis. Biol. Bull. 182, 169–176 (1992).

25. Steger, J. et al. Single-cell transcriptomics identifies conserved regulators of neuroglandular lineages. Cell Rep. 40, (2022).

26. Cole, A. G. et al. Updated single cell reference atlas for the starlet anemone Nematostella vectensis. Front. Zool. 21, 8 (2024).

27. Siebert, S. et al. Stem cell differentiation trajectories in Hydra resolved at single-cell resolution. Science 365, eaav9314 (2019).

28. Chari, T. et al. Whole-animal multiplexed single-cell RNA-seq reveals transcriptional shifts across Clytia medusa cell types. Sci. Adv. 7, eabh1683.

29. Hu, M., Zheng, X., Fan, C.-M. & Zheng, Y. Lineage dynamics of the endosymbiotic cell type in the soft coral Xenia. Nature 582, 534–538 (2020).

30. Levy, S. et al. A stony coral cell atlas illuminates the molecular and cellular basis of coral symbiosis, calcification, and immunity. Cell 184, 2973–2987.e18 (2021).

31. Levy, S. et al. The evolution of facultative symbiosis in stony corals. Nature 1–9 (2025) doi:10.1038/s41586-025-09623-6.

32. Farrell, J. A. et al. Single-cell reconstruction of developmental trajectories during zebrafish embryogenesis. Science 360, eaar3131 (2018).

33. Layden, M. J., Boekhout, M. & Martindale, M. Q. Nematostella vectensis achaete-scute homolog NvashA regulates embryonic ectodermal neurogenesis and represents an ancient component of the metazoan neural specification pathway. Dev. Camb. Engl. 139, 1013–1022 (2012).

34. Richards, G. S. & Rentzsch, F. Transgenic analysis of a SoxB gene reveals neural progenitor cells in the cnidarian Nematostella vectensis. Development 141, 4681–4689 (2014).

35. Takahashi, T. Comparative Aspects of Structure and Function of Cnidarian Neuropeptides. Front. Endocrinol. 11, (2020).

36. Tournière, O., Gahan, J. M., Busengdal, H., Bartsch, N. & Rentzsch, F. Insm1-expressing neurons and secretory cells develop from a common pool of progenitors in the sea anemone Nematostella vectensis. Sci. Adv. 8, eabi7109.

37. Piovani, L. et al. Single-cell atlases of two lophotrochozoan larvae highlight their complex evolutionary histories. Sci. Adv. 9, eadg6034 (2023).

38. Barrera-Redondo, J., Lotharukpong, J. S., Drost, H.-G. & Coelho, S. M. Uncovering gene-family founder events during major evolutionary transitions in animals, plants and fungi using GenEra. Genome Biol. 24, 54 (2023).

39. Drost, H.-G., Gabel, A., Liu, J., Quint, M. & Grosse, I. myTAI: evolutionary transcriptomics with R. Bioinformatics 34, 1589–1590 (2018).

40. David, C. N. et al. Evolution of complex structures: minicollagens shape the cnidarian nematocyst. Trends Genet. 24, 431–438 (2008).

41. Peters, E. C. et al. NOMENCLATURE AND BIOLOGY OF ASTRANGIA POCULATA i=A. DANAE, =A. ASTREIFORMIS).

42. Zenkert, C., Takahashi, T., Diesner, M.-O. & Özbek, S. Morphological and Molecular Analysis of the Nematostella vectensis Cnidom. PLoS ONE 6, e22725 (2011).

43. Babonis, et al., L. S. A novel regulatory gene promotes novel cell fate by suppressing ancestral fate in the sea anemone Nematostella vectensis. https://www.pnas.org/doi/10.1073/pnas.2113701119 (2022) doi:10.1073/pnas.2113701119.

44. Babonis, L. S. & Martindale, M. Q. PaxA, but not PaxC, is required for cnidocyte development in the sea anemone Nematostella vectensis. EvoDevo 8, 14 (2017).

45. Östman, C. A guideline to nematocyst nomenclature and classification, and some notes on the systematic value of nematocysts. Sci. Mar. 64, 31–46 (2000).

46. Plessier, F. & Marlow, H. Cellular and transcriptional trajectories of neural fate specification in sea anemone uncover two modes of adult neurogenesis. Nat. Commun. 17, 2611 (2026).

47. Karabulut, A., He, S., Chen, C.-Y., McKinney, S. A. & Gibson, M. C. Electroporation of short hairpin RNAs for rapid and efficient gene knockdown in the starlet sea anemone, *Nematostella vectensis*. Dev. Biol. 448, 7–15 (2019).

48. Shafer, M. E. R. Cross-Species Analysis of Single-Cell Transcriptomic Data. Front. Cell Dev. Biol. 7, (2019).

49. Song, Y., Miao, Z., Brazma, A. & Papatheodorou, I. Benchmarking strategies for cross-species integration of single-cell RNA sequencing data. Nat. Commun. 14, 6495 (2023).

50. True, J. R. & Haag, E. S. Developmental system drift and flexibility in evolutionary trajectories. Evol. Dev. 3, 109–119 (2001).

51. McColgan, Á. & DiFrisco, J. Understanding developmental system drift. Development 151, dev203054 (2024).

52. Ashey, J., Putnam, H. M. & McManus, M. C. Guided by the northern star coral: a research synthesis and roadmap for Astrangia poculata. Biol. Lett. 21, 20240469 (2025).

53. Stankiewicz, K. H. et al. Genomic comparison of the temperate coral Astrangia poculata with tropical corals yields insights into winter quiescence, innate immunity, and sexual reproduction. G3 GenesGenomesGenetics jkaf033 (2025) doi:10.1093/g3journal/jkaf033.

54. Stefanik, D. J., Friedman, L. E. & Finnerty, J. R. Collecting, rearing, spawning and inducing regeneration of the starlet sea anemone, Nematostella vectensis. Nat. Protoc. 8, 916–923 (2013).

55. Fritzenwanker, J. H. & Technau, U. Induction of gametogenesis in the basal cnidarian Nematostella vectensis (Anthozoa). Dev. Genes Evol. 212, 99–103 (2002).

56. Babonis, L. S., Martindale, M. Q. & Ryan, J. F. Do novel genes drive morphological novelty? An investigation of the nematosomes in the sea anemone Nematostella vectensis. BMC Evol. Biol. 16, 114 (2016).

57. Nakanishi, N. & Martindale, M. Q. CRISPR knockouts reveal an endogenous role for ancient neuropeptides in regulating developmental timing in a sea anemone. eLife 7, e39742 (2018).

58. Oren, M., Brikner, I., Appelbaum, L. & Levy, O. Fast Neurotransmission Related Genes Are Expressed in Non Nervous Endoderm in the Sea Anemone Nematostella vectensis. PLOS ONE 9, e93832 (2014).

59. Schindelin, J., et al. Fiji: an open-source platform for biological-image analysis. Nat. Methods 9, 676–682 (2012).

60. Wolenski, F. S., Layden, M. J., Martindale, M. Q., Gilmore, T. D. & Finnerty, J. R. Characterizing the spatiotemporal expression of RNAs and proteins in the starlet sea anemone, Nematostella vectensis. Nat. Protoc. 8, 900–915 (2013).

61. Shpirer, E. et al. Diversity and evolution of myxozoan minicollagens and nematogalectins. BMC Evol. Biol. 14, 205 (2014).

62. Katoh, K. & Standley, D. M. MAFFT Multiple Sequence Alignment Software Version 7: Improvements in Performance and Usability. Mol. Biol. Evol. 30, 772–780 (2013).

63. Stamatakis, A. RAxML version 8: a tool for phylogenetic analysis and post-analysis of large phylogenies. Bioinformatics 30, 1312–1313 (2014).

64. Wong, T. K. F. et al. IQ-TREE 3: phylogenomic inference software using complex evolutionary models. Mol. Biol. Evol. 43, msag117 (2026).

65. Rambaut, A. FigTree v1.3.1. https://tree.bio.ed.ac.uk/software/figtree/ (2010).

66. Wickham, H. Getting Started with ggplot2. in ggplot2: Elegant Graphics for Data Analysis (ed. Wickham, H.) 11–31 (Springer International Publishing, Cham, 2016). doi:10.1007/978-3-319-24277-4_2.

67. Massri, A. J. et al. Developmental single-cell transcriptomics in the Lytechinus variegatus sea urchin embryo. Development 148, dev198614 (2021).

68. Zheng, G. X. Y. et al. Massively parallel digital transcriptional profiling of single cells. Nat. Commun. 8, 14049 (2017).

69. Stanke, M., Diekhans, M., Baertsch, R. & Haussler, D. Using native and syntenically mapped cDNA alignments to improve *de novo* gene finding. Bioinformatics 24, 637–644 (2008).

70. Li, H. et al. The Sequence Alignment/Map format and SAMtools. Bioinformatics 25, 2078–2079 (2009).

71. Hoff, K. J. & Stanke, M. Predicting Genes in Single Genomes with AUGUSTUS. Curr. Protoc. Bioinforma. 65, e57 (2019).

72. Keilwagen, J., Hartung, F. & Grau, J. GeMoMa: Homology-Based Gene Prediction Utilizing Intron Position Conservation and RNA-seq Data. Methods Mol. Biol. 1962, 161–177 (2019).

73. Keilwagen, J. et al. Using intron position conservation for homology-based gene prediction. Nucleic Acids Res. 44, e89 (2016).

74. Keilwagen, J., Hartung, F., Paulini, M., Twardziok, S. O. & Grau, J. Combining RNA-seq data and homology-based gene prediction for plants, animals and fungi. BMC Bioinformatics 19, 189 (2018).

75. Hao, Y. et al. Dictionary learning for integrative, multimodal and scalable single-cell analysis. Nat. Biotechnol. 42, 293–304 (2024).

76. Virshup, I., Rybakov, S., Theis, F. J., Angerer, P. & Wolf, F. A. anndata: Access and store annotated data matrices. J. Open Source Softw. 9, 4371 (2024).

77. Kolde, R. pheatmap: Pretty Heatmaps. R-universe https://raivokolde.r-universe.dev/pheatmap (2025).

78. Gu, Z. Complex heatmap visualization. iMeta 1, e43 (2022).

79. Gu, Z., Eils, R. & Schlesner, M. Complex heatmaps reveal patterns and correlations in multidimensional genomic data. Bioinformatics 32, 2847–2849 (2016).

80. Allaire, J. J., Gandrud, C., Russell, K. & Yetman, C. networkD3: D3 JavaScript Network Graphs from R. https://github.com/christophergandrud/networkd3 (2025).

81. Cantalapiedra, C. P., Hernández-Plaza, A., Letunic, I., Bork, P. & Huerta-Cepas, J. eggNOG-mapper v2: Functional Annotation, Orthology Assignments, and Domain Prediction at the Metagenomic Scale. Mol. Biol. Evol. 38, 5825–5829 (2021).

